# Identification of the bacteriophage nucleus protein interaction network

**DOI:** 10.1101/2023.05.18.541317

**Authors:** Eray Enustun, Amar Deep, Yajie Gu, Katrina T. Nguyen, Vorrapon Chaikeeratisak, Emily Armbruster, Majid Ghassemian, Elizabeth Villa, Joe Pogliano, Kevin D. Corbett

## Abstract

In the arms race between bacteria and bacteriophages (phages), some large-genome jumbo phages have evolved a protein shell that encloses their replicating genome to protect it against DNA-targeting immune factors. By segregating the genome from the host cytoplasm, however, the “phage nucleus” introduces the need to specifically transport mRNA and proteins through the nuclear shell, and to dock capsids on the shell for genome packaging. Here, we use proximity labeling and localization mapping to systematically identify proteins associated with the major nuclear shell protein chimallin (ChmA) and other distinctive structures assembled by these phages. We identify six uncharacterized nuclear shell-associated proteins, one of which directly interacts with self-assembled ChmA. The structure and protein-protein interaction network of this protein, which we term ChmB, suggests that it forms pores in the ChmA lattice that serve as docking sites for capsid genome packaging, and may also participate in mRNA and/or protein transport.

## Introduction

Since the discovery of phages more than a century ago, research on these remarkable organisms has yielded fundamental insights into a broad range of pathways across biology^1^. Historically, most phage studies have focused on small-genome phages (∼30-140 kb), leaving larger “jumbo phages” with genomes over ∼200 kb much less well understood despite their abundance in nature^2–4^. We previously showed that one family of jumbo phages forms distinctive structures in infected cells, including a nucleus-like compartment bounded by a proteinaceous shell and a spindle-like structure that centers and rotates this compartment^5–7^. The phage nucleus encloses the replicating phage genome and excludes most host proteins including CRISPR effectors and restriction enzymes, rendering this family of phages broadly resistant to DNA-targeting bacterial immune systems^8, 9^.

The nuclear shell provides an important selective advantage to the phage by protecting its replicating genome from host-encoded defense nucleases, but that protection comes at the cost of significant added complexity in the phage life cycle. The phage nuclear shell is composed mostly of one protein, termed chimallin (ChmA), which forms a single-layer thick flexible lattice that separates the phage genome from the bacterial cytoplasm^10, 11^. Pores in the ChmA lattice are less than ∼2 nm in width, large enough to pass metabolites but too small for passage of most proteins or mRNAs^10^. As in eukaryotes, mRNAs are transcribed within the phage nucleus but translated in the cytoplasm, meaning that phage mRNAs must be transported out of the nucleus^6^. At the same time, phage-encoded proteins necessary for genome replication and mRNA transcription must be specifically transported back into the nucleus^6, 12^. Finally, during virion production newly assembled capsids are trafficked along filaments of the tubulin-like protein PhuZ to the nuclear shell, where they dock for genome packaging^6, 13^. In this family of phages, final assembly of capsids with virion tails occurs at a pair of structures termed the “phage bouquets” prior to cell lysis and virion release^14, 15^.

The diverse functions of the jumbo phage nuclear shell – including mRNA export, protein import, and capsid docking – imply that this structure incorporates multiple components in addition to the ChmA shell protein that mediate these functions. Here, we use proximity labeling (miniTurboID^16^) in *Pseudomonas aeruginosa* cells infected by the jumbo phage ΦPA3^17^ to identify proteins that localize both within the phage nucleus and specifically to the nuclear shell. We identify six new nuclear shell-associated proteins, one of which interacts directly with both ChmA and the putative portal protein^18^. These interactions suggest that this protein, which we term ChmB, forms pores in the ChmA lattice and mediates capsid docking and genome packaging. ChmB’s overall protein-protein interaction network further suggests additional roles in mRNA export and/or protein import into the phage nucleus. More broadly, our data define a jumbo phage nuclear shell interaction network and reveal the subcellular localization of dozens of previously uncharacterized jumbo phage proteins.

## Results

### Identification of jumbo phage nucleus-associated proteins

The genomes of nucleus-forming jumbo phages are poorly characterized: for instance, 290 of the 378 genes encoded by the *Pseudomonas* jumbo phage ΦPA3 have no annotated function in the NCBI protein database. To overcome this deficit, we used a proximity labeling approach to identify proteins associated with the phage nuclear shell that could endow this structure with additional functionality like mRNA export, protein import, and capsid docking. We fused the promiscuous biotin ligase miniTurboID^16^ to the ChmA protein from the jumbo phage ΦPA3 (gp53) and to the phage’s nuclear-localized RecA protein (gp175) (**Figure 1a-b, S1a-b**)7. We expressed each of these fusion proteins in *Pseudomonas aeruginosa* cells infected with ΦPA3, collected samples at 45 minutes post infection (mpi) which is the earliest time point when the mature nuclear shell was observed with docked capsids^7^, and performed streptavidin pulldown and mass spectrometry analysis to identify biotinylated proteins. By focusing on the phage proteins that were biotinylated and normalizing the results to a control miniTurboID-GFP fusion (which remains diffuse in the host cell cytoplasm throughout infection; **Figure S1b**), we identified candidate proteins that preferentially localize in close proximity to RecA and/or ChmA (**Tables 1-2, Supplemental Tables 1-2**).

**Figure 1.**
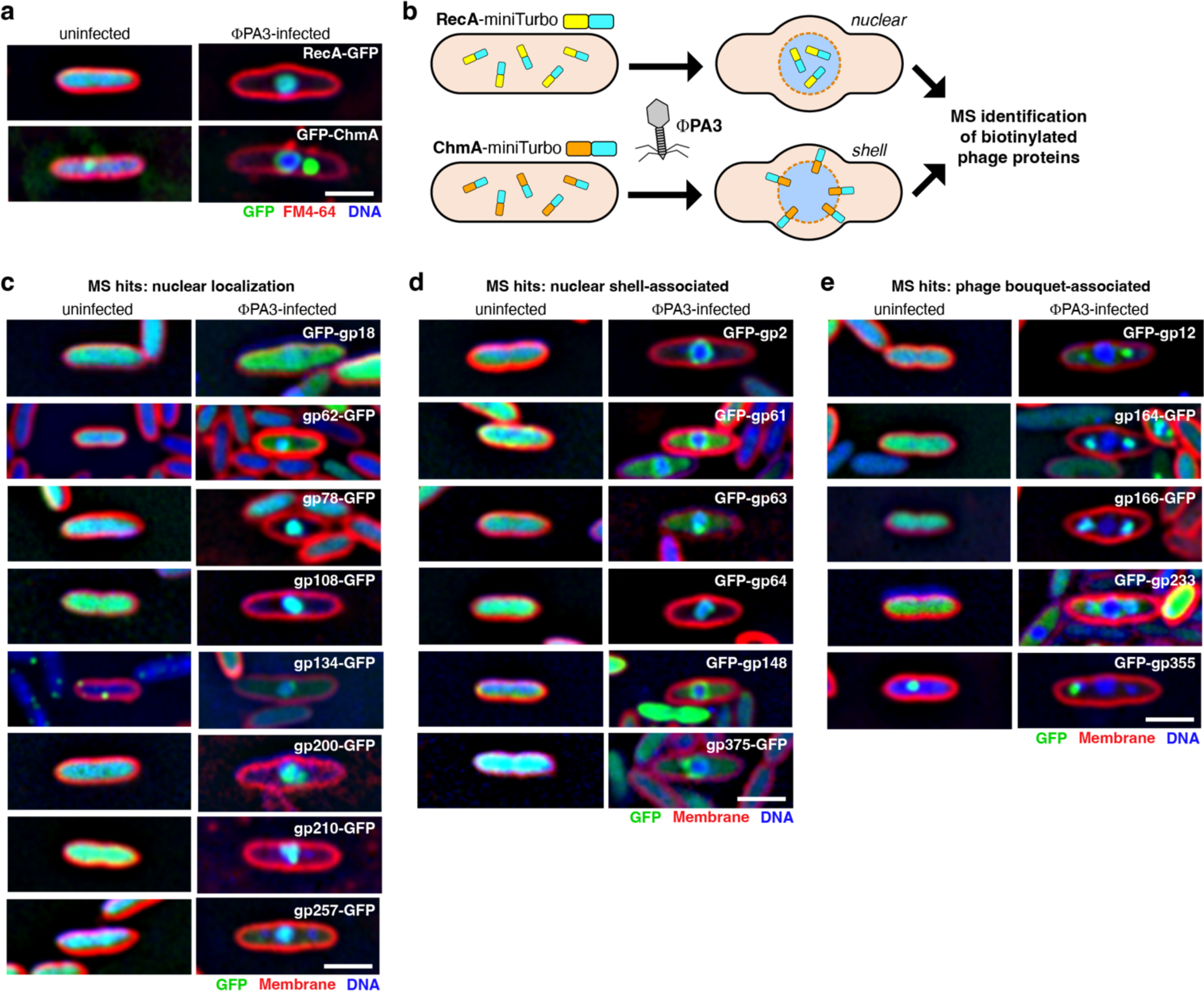
Identification of jumbo phage nuclear shell-associated proteins. (a) Subcellular localization of GFP-tagged ΦPA3 RecA (gp175) and ChmA (gp53) in uninfected (left) and ΦPA3-infected (right) *P. aeruginosa* cells. GFP is shown in green, FM4-64 (to visualize membranes) in red, and DAPI (to visualize nucleic acids) in blue. Scale bar = 2 µm. (b) Experimental schematic for identification of jumbo phage nuclear or nuclear-shell-associated genes, by proximity labeling with miniTurboID-fused RecA or ChmA in ΦPA3-infected *P. aeruginosa* cells. See **Figure S1a** for fusion construct design, and **Figure S1b** for localization of miniTurboID-fused proteins. See **Tables 1** and **2** for top 25 identified proteins, **Tables S1** and **S2** for full protein lists, and **Figure S1c-d** for diagrams showing overlap between independent mass spectrometry datasets. (c-e) Subcellular localization of selected proteins identified by proximity labeling, with panel (c) showing nuclear-localized proteins, (d) showing nuclear shell-associated proteins, and (e) showing phage bouquet-associated proteins. See **Figure S2** for further data, and **Table 3** for a collated list of localizations. Scale bar = 2 µm.

**Table 1.**
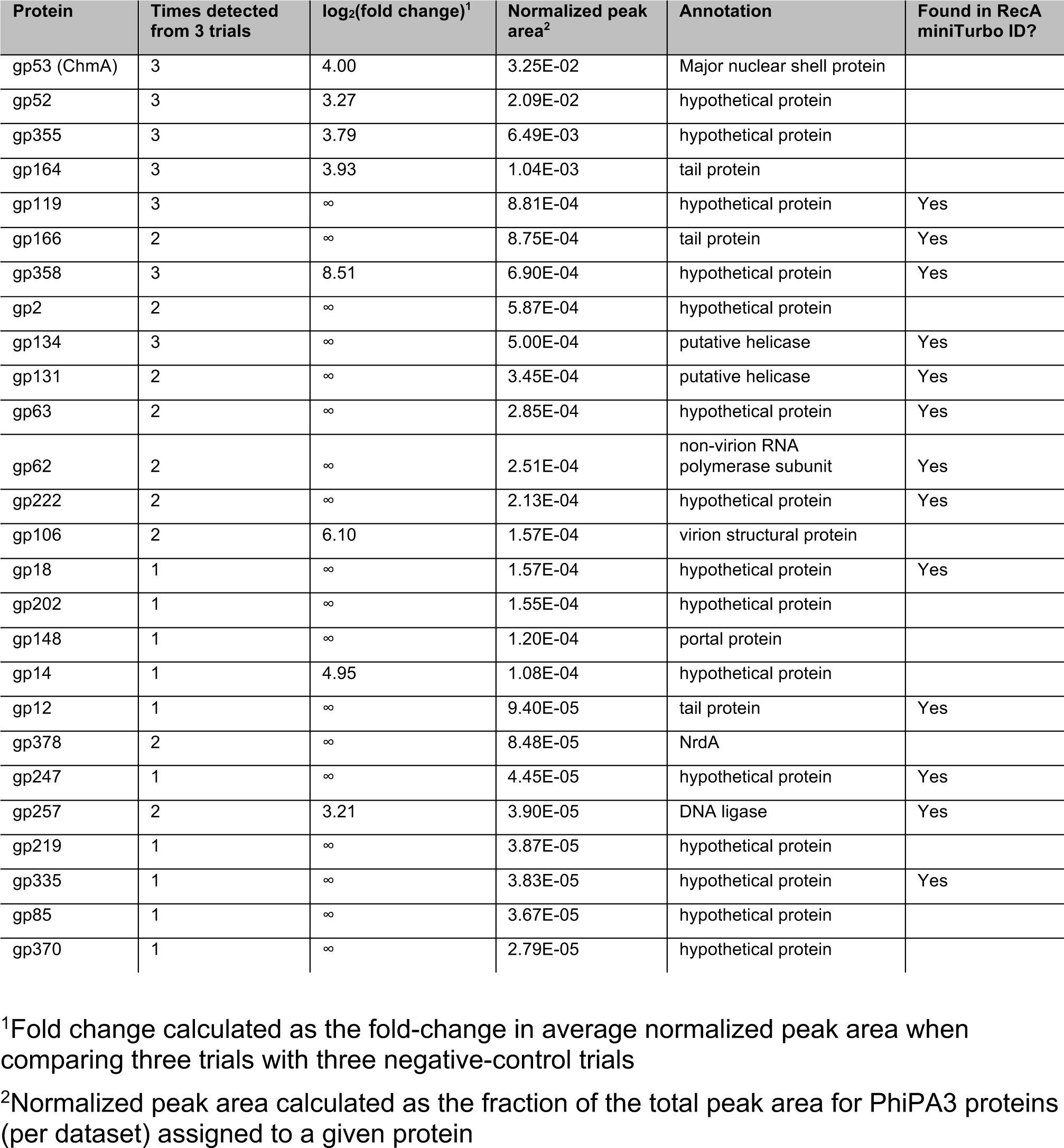
Top 25 identified proteins from ΦPA3 ChmA (gp53) miniTurboID.

**Table 2.**
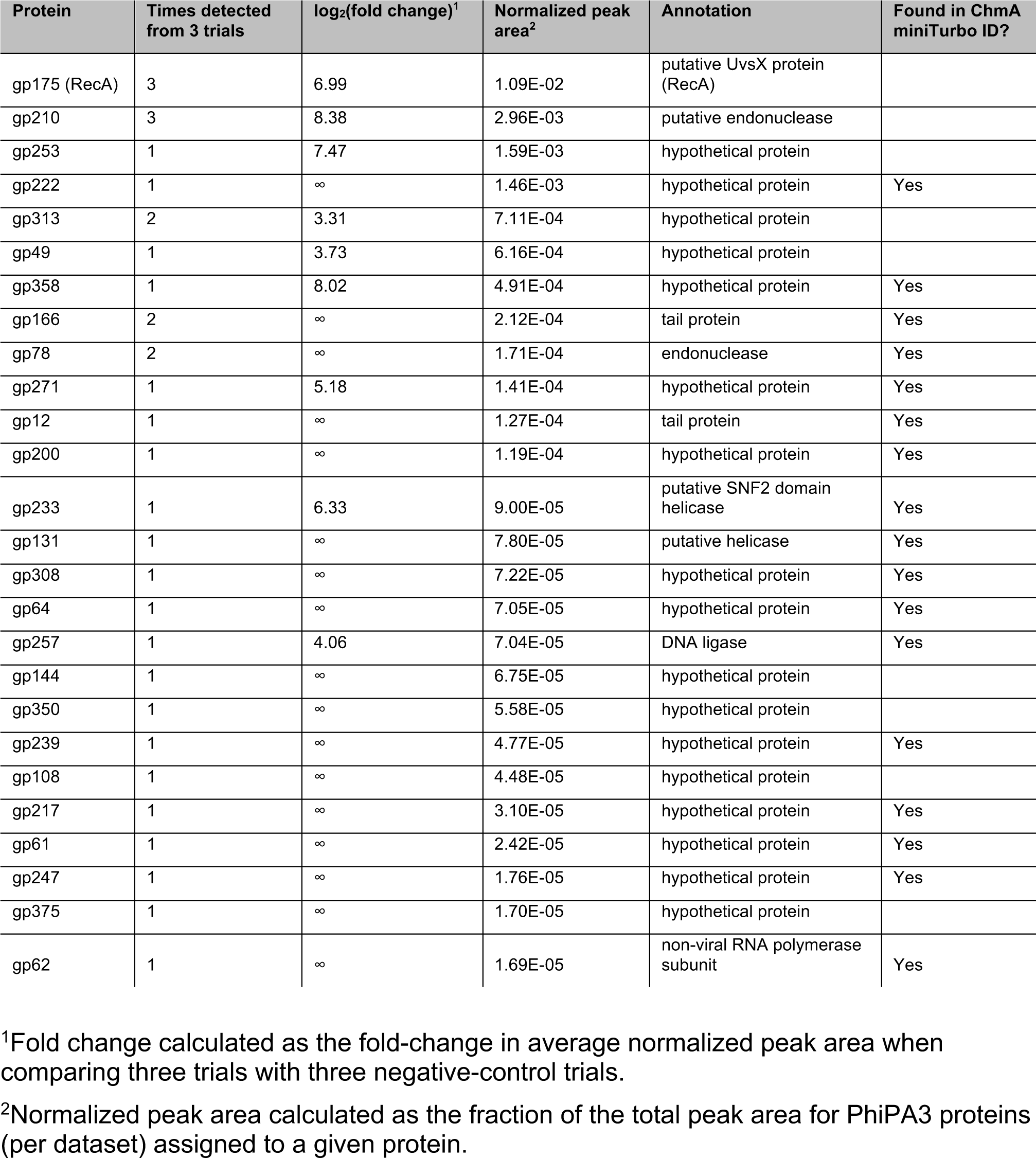
Top 25 identified proteins from ΦPA3 RecA (gp175) miniTurboID.

| Protein | Times detected from 3 trials | $\log_2(\text{fold change})^1$ | Normalized peak area<sup>2</sup> | Annotation | Found in ChmA miniTurbo ID? |
| --- | --- | --- | --- | --- | --- |
| gp175 (RecA) | 3 | 6.99 | 1.09E-02 | putative UvsX protein (RecA) |  |
| gp210 | 3 | 8.38 | 2.96E-03 | putative endonuclease |  |
| gp253 | 1 | 7.47 | 1.59E-03 | hypothetical protein |  |
| gp222 | 1 | ∞ | 1.46E-03 | hypothetical protein | Yes |
| gp313 | 2 | 3.31 | 7.11E-04 | hypothetical protein |  |
| gp49 | 1 | 3.73 | 6.16E-04 | hypothetical protein |  |
| gp358 | 1 | 8.02 | 4.91E-04 | hypothetical protein | Yes |
| gp166 | 2 | ∞ | 2.12E-04 | tail protein | Yes |
| gp78 | 2 | ∞ | 1.71E-04 | endonuclease | Yes |
| gp271 | 1 | 5.18 | 1.41E-04 | hypothetical protein | Yes |
| gp12 | 1 | ∞ | 1.27E-04 | tail protein | Yes |
| gp200 | 1 | ∞ | 1.19E-04 | hypothetical protein | Yes |
| gp233 | 1 | 6.33 | 9.00E-05 | putative SNF2 domain helicase | Yes |
| gp131 | 1 | ∞ | 7.80E-05 | putative helicase | Yes |
| gp308 | 1 | ∞ | 7.22E-05 | hypothetical protein | Yes |
| gp64 | 1 | ∞ | 7.05E-05 | hypothetical protein | Yes |
| gp257 | 1 | 4.06 | 7.04E-05 | DNA ligase | Yes |
| gp144 | 1 | ∞ | 6.75E-05 | hypothetical protein | |
| gp350 | 1 | ∞ | 5.58E-05 | hypothetical protein | |
| gp239 | 1 | ∞ | 4.77E-05 | hypothetical protein | Yes |
| gp108 | 1 | ∞ | 4.48E-05 | hypothetical protein | |
| gp217 | 1 | ∞ | 3.10E-05 | hypothetical protein | Yes |
| gp61 | 1 | ∞ | 2.42E-05 | hypothetical protein | Yes |
| gp247 | 1 | ∞ | 1.76E-05 | hypothetical protein | Yes |
| gp375 | 1 | ∞ | 1.70E-05 | hypothetical protein | |
| gp62 | 1 | ∞ | 1.69E-05 | non-viral RNA polymerase subunit | Yes |

<sup>1</sup>Fold change calculated as the fold-change in average normalized peak area when comparing three trials with three negative-control trials.
<sup>2</sup>Normalized peak area calculated as the fraction of the total peak area for ΦPA3 proteins (per dataset) assigned to a given protein.

**Table 3.** Localization of miniTurboID hits in infected cells.

| Protein | Annotation | Detected with ChmA or RecA? | Localization in infected cells |
| --- | --- | --- | --- |
| gp2 | hypothetical protein | ChmA | shell |
| gp12 | tail protein | both | phage bouquet |
| gp14 | hypothetical protein | ChmA | cytoplasmic puncta |
| gp18 | hypothetical protein | both | nuclear |
| gp49 | hypothetical protein | RecA | diffuse cytoplasmic |
| gp52 | hypothetical protein | ChmA | cytoplasmic puncta |
| gp53 | Major nuclear shell protein (ChmA) | ChmA | shell |
| gp61 | hypothetical protein | both | shell |
| gp62 | non-virion RNA polymerase | both | nuclear |
| gp63 | hypothetical protein | both | shell |
| gp64 | hypothetical protein | both | shell |
| gp78 | endonuclease | both | nuclear |
| gp85 | hypothetical protein | ChmA | diffuse cytoplasmic |
| gp106 | virion structural protein | ChmA | cytoplasmic puncta |
| gp108 | hypothetical protein | RecA | nuclear |
| gp119 | hypothetical protein | both | cytoplasmic puncta |
| gp131 | putative helicase | both | cytoplasmic puncta |
| gp134 | putative helicase | both | nuclear |
| gp144 | hypothetical protein | RecA | cytoplasmic |
| gp148 | portal protein | ChmA | shell |
| gp164 | tail protein | ChmA | phage bouquet |
| gp166 | tail protein | both | phage bouquet |
| gp175 | putative UvsX protein (RecA) | RecA | nuclear |
| gp200 | hypothetical protein | both | nuclear |
| gp202 | hypothetical protein | ChmA | diffuse cytoplasmic |
| gp210 | putative endonuclease | RecA | nuclear |
| gp217 | hypothetical protein | both | cytoplasmic puncta |
| gp219 | hypothetical protein | ChmA | cytoplasmic puncta |
| gp222 | hypothetical protein | both | diffuse cytoplasmic |
| gp233 | putative SNF2 domain helicase | both | phage bouquet |
| gp239 | hypothetical protein | both | diffuse cytoplasmic |
| gp247 | hypothetical protein | both | diffuse cytoplasmic |
| gp253 | hypothetical protein | RecA | cytoplasmic puncta |
| gp257 | DNA ligase | both | nuclear |
| gp271 | hypothetical protein | RecA | diffuse cytoplasmic |
| gp308 | hypothetical protein | both | cytoplasmic puncta |
| gp313 | hypothetical protein | RecA | diffuse cytoplasmic |
| gp335 | hypothetical protein | both | diffuse cytoplasmic |
| gp350 | hypothetical protein | RecA | diffuse cytoplasmic |
| gp355 | hypothetical protein | ChmA | phage bouquet/cytoplasmic puncta |
| gp358 | hypothetical protein | both | cytoplasmic puncta |
| gp370 | hypothetical protein | ChmA | diffuse cytoplasmic |
| gp375 | hypothetical protein | RecA | shell |
| gp378 | NrdA | ChmA | diffuse cytoplasmic |

To validate our interaction data, we generated GFP fusions of the top 25 ChmA- and RecA-interacting proteins from our miniTurboID datasets (42 proteins total plus ChmA and RecA; **Table 3**), expressed each in *P. aeruginosa*, and determined their localization in both uninfected and ΦPA3-infected cells. Proteins that interact with RecA are expected to localize inside the phage nucleus, while ChmA-interacting proteins are expected to localize on or near the nuclear shell. Given the architecture of the ChmA lattice, we expect the N-terminal miniTurboID tag to be localized on the outer surface of the nuclear shell^10^. Among the 42 proteins tested, we identified eight that localize within the nucleus-like compartment, six that localize to the nuclear shell itself, and five that localize to the phage bouquets (**Figure 1c-e**). Most proteins that we find localizes within the nucleus are annotated in the NCBI database to be interacting with nucleic acids, including a subunit of the phage-encoded non-virion RNA polymerase (nvRNAP; gp62)^19^, two predicted helicases (gp131 and gp134), a predicted DNA ligase (gp257), and two predicted endonucleases (gp78 and gp210) (**Figure 1c, Table 3**)^18^. Two nuclear-localized proteins (gp108 and gp200) have no annotated or predicted function. Of the five proteins that localize to phage bouquets, three (gp12, gp164, and gp166) are predicted phage tail proteins, one (gp233) is a predicted helicase, and one (gp355) has no annotated or predicted function^18, 20, 21^.

To date, the only known component of the jumbo phage nuclear shell is ChmA^6, 7, 10^. Among our list of RecA- and ChmA-interacting proteins, we identified six proteins (gp2, gp61, gp63, gp64, gp148, and gp375) that clearly localize to the nuclear shell upon ΦPA3 infection of *P. aeruginosa* cells (**Figure 1d, Table 3**). One of these proteins, gp148, its homolog was previously predicted to be the portal protein of the phage capsid^18^. In other phages, the portal protein forms a homododecameric complex that orchestrates capsid assembly^22^ and associates with the terminase to translocate genomic DNA into the empty capsid^23–27^. We previously showed that in this family of jumbo phages, capsids are docked on the nuclear shell for genomic DNA packaging^6^, and our finding that this putative portal protein associates with the shell suggests that it is directly responsible for capsid docking. Notably, we find that gp148 localizes to the phage nuclear shell as early as 30 minutes post infection, well before capsid assembly and docking begins at around 45 minutes post infection^6^ (**Figure S3**). This finding suggests that the portal protein can localize to the nuclear shell on its own, supporting the idea that it directly mediates capsid docking.

Apart from the portal protein, the remaining five shell-localized proteins have no predicted function and were not found in prior mass spectrometry studies of fully formed virions^20^. All five proteins are conserved across jumbo phages (gp61, gp63, gp64) or within *Pseudomonas*-infecting jumbo phages (gp2 and gp375), but show no detectable sequence homology to any other known proteins. Moreover, all five proteins are expressed early in infections, with timing similar to the major nuclear shell protein ChmA^6, 7^. Thus, we speculate that some or all of these proteins are components of the nuclear shell itself that may mediate mRNA export, protein import, and/or capsid docking.

### gp2 is an interaction hub at the nuclear shell

Among the identified nuclear shell-associated proteins, gp2 was one of the most highly biotinylated proteins in our miniTurboID-ChmA samples (**Table 1**). We verified that GFP fusions of both ΦPA3 gp2 and its homolog from the related jumbo phage 201Φ2-1 (gp2) colocalize with mCherry-fused ChmA in infected cells (**Figure 2a,b**). To further define the interaction network of ΦPA3 gp2, we expressed GFP-fused gp2 in ΦPA3-infected *P. aeruginosa* cells, then purified the protein and interacting partners using GFP affinity chromatography. After purification of gp2 with a C-terminal GFP tag, we identified two strong bands at a molecular weight of ∼70 kDa on a silver-stained SDS-PAGE gel (**Figure 2c**). We extracted a gel slice containing these two closely spaced bands and used trypsin mass spectrometry to identify the proteins. In this sample, the strongest signal (149 peptides, 81% sequence coverage) was for ChmA (gp53, 66.7 kDa), and the second-strongest (41 peptides, 40% sequence coverage) was for the major capsid protein (gp136, 82.8 kDa; **Table S3**). These data suggest that the observed doublet at ∼70 kDa represents these two proteins, and that gp2 interacts strongly with both ChmA and capsids.

**Figure 2.**
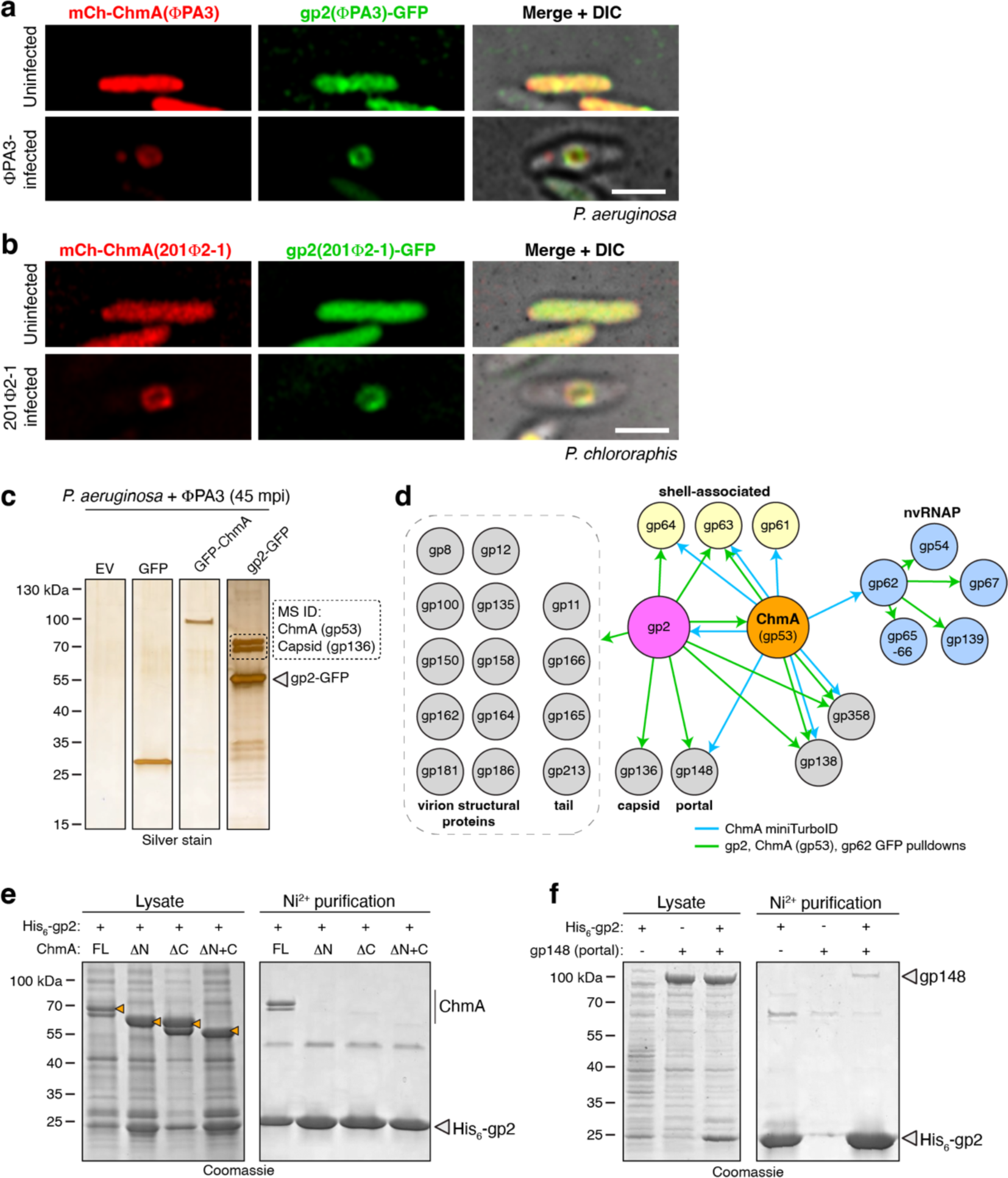
gp2 is an interaction hub in the jumbo phage nuclear shell. (a) Colocalization of mCherry-fused ΦPA3 ChmA (red) and GFP-fused ΦPA3 gp2 (green) in *P. aeruginosa* cells. Scale bar = 2 µm. (b) Colocalization of mCherry-fused 201Φ2-1 ChmA (gp105; red) and GFP-fused 201Φ2-1 gp2 (green) in *P. chlororaphis* cells. Scale bar = 2 µm. (c) Silver stain SDS-PAGE analysis of GFP pulldown experiments. EV: empty vector. Dotted box indicated the gel slice that was cut out (of the same bands in a Coomassie blue-stained gel) for tryptic mass spectrometry protein identification (see **Table S3**). (d) Interaction network of the jumbo phage nuclear shell, with blue arrows indicating interactions identified by ChmA miniTurboID and green arrows indicating interactions identified in GFP pulldowns (see **Figure S4a** for SDS-PAGE gels of all analyzed GFP pulldown samples, and **Table S4** for full data). nvRNAP: non-virion RNA polymerase. (e) Ni^2+^ pulldown analysis of *E. coli*-coexpressed 201Φ2-1 gp2 (His6-tagged) and ChmA (full-length or truncated: ΔN missing residues 1-63, ΔC missing residues 583-631, and ΔN+C missing residues 1-63 and 583-631). Doublet bands for ChmA arise from a methionine codon at position 33 of the annotated gene. Orange marks show the presence of ChmA in the lysates. See **Figure S4b** for control pulldown. (f) Ni^2+^ pulldown analysis of *E. coli*-coexpressed ΦPA3 gp2 (His6-tagged) and portal (gp148).

To further investigate the interactions between nuclear shell-associated proteins, we next performed mass spectrometry on the full purified samples from GFP-tagged ChmA, gp2 (both N- and C-terminal GFP tags), and gp62 (**Figure 2d**, **Figure S4a**, **Table S4**). We confirmed that this approach is successful in purifying functional protein complexes, as gp62 is predicted to be part of a five-subunit non-virion RNA polymerase^19^, and mass spectrometry successfully identified all other subunits of this complex in the gp62-GFP pulldown (**Figure 2d**, **Table S4**). The most enriched protein in GFP-tagged gp2 samples was ChmA, and we also identified the major capsid protein (gp136), the predicted portal protein (gp148), and 14 other proteins annotated as either tail or virion structural proteins (**Figure 2d**, **Table S4**). Some of these proteins were also identified in pulldowns with other GFP-tagged proteins, suggesting that the proteins may simply be highly abundant in cell lysates (**Table S4**). Nonetheless, the strong enrichment of phage structural proteins in GFP-tagged gp2 pulldowns strongly suggest that gp2 interacts directly with assembling virions, potentially as they dock on the nuclear shell for filling with genomic DNA. Also identified in the GFP-tagged gp2 pulldowns were two other shell-associated proteins, gp63 and gp64 (**Figure 2d**, **Table S4**), suggesting that these proteins may interact either directly with gp2, or instead interact indirectly through the ChmA lattice. Finally, two additional proteins (gp138 and gp358) were identified in both the ChmA and gp2 GFP pulldowns, as well as having been detected by ChmA-miniTurboID labeling (**Figure 2d**, **Table S1**, **Table S4**). These two proteins are conserved across jumbo phages infecting *Pseudomonas* but have no annotated or predicted function, leaving their potential roles unknown.

### gp2 interacts directly with ChmA and the phage portal protein

We previously showed that ChmA from two jumbo phages, 201Φ2-1 and Goslar, self-assembles into closed structures resembling the phage nuclear shell when expressed in *E. coli*^10^. To determine whether gp2 interacts directly with ChmA, we co-expressed 201Φ2-1 gp2 and ChmA in *E. coli* and performed Ni^2+^ pulldowns using a His_6_-tag on gp2. We found that His_6_-gp2 robustly interacts with ChmA in this assay, showing that the two proteins directly interact (**Figure 2e, S4b**). We next deleted the N- and C-terminal segments (NTS and CTS, respectively), which bind to neighboring protomers in the ChmA lattice to mediate nuclear shell assembly. In vitro, deletion of either NTS or CTS leads to a loss of ChmA self-assembly^10^, and we found that gp2 was unable to interact with ChmA mutants lacking NTS, CTS, or both NTS and CTS (**Figure 2e**)^10^. The loss of gp2 binding when deleting either the ChmA NTS or CTS suggests that gp2 does not simply bind one of these tail segments; rather the data suggest that gp2 interacts specifically with the assembled ChmA lattice.

We next performed a similar coexpression experiment with His_6_-tagged ΦPA3 gp2 and the portal protein gp148. We found that gp2 was able to interact directly with gp148 in this assay (**Figure 2f**). Combined with our data showing that gp2 interacts directly with self-assembled ChmA, these data suggest that gp2 is an integral component of the phage nuclear shell that is directly involved in the docking and filling of capsids through an interaction with the portal. Thus, we name ΦPA3 gp2 and its homologs in related jumbo phages chimallin B (ChmB).

### ChmB forms a homodimer with a novel fold

To determine the structural basis for ChmB interactions with other proteins, we recombinantly purified the protein from several jumbo phages and determined a 2.6 Å resolution crystal structure of ChmB from the related phage PA1C (gp2; 38% identical to ΦPA3 gp2) (**Table S5**). ChmB forms a homodimer in solution (**Figure 3a, S4A-C**), and the structure reveals a distinctive intertwined dimer with each protomer’s N-terminus forming a short ꞵ-strand and an ɑ-helix that pack against the C-terminal globular domain of its dimer mate. Overall, the ChmB dimer adopts a distinctive U shape with dimensions of ∼5 × 8 nm (**Figure 3b**). Searches with the DALI or FoldSeek protein structure comparison tools^28, 29^ show no known structural relatives.

**Figure 3.**
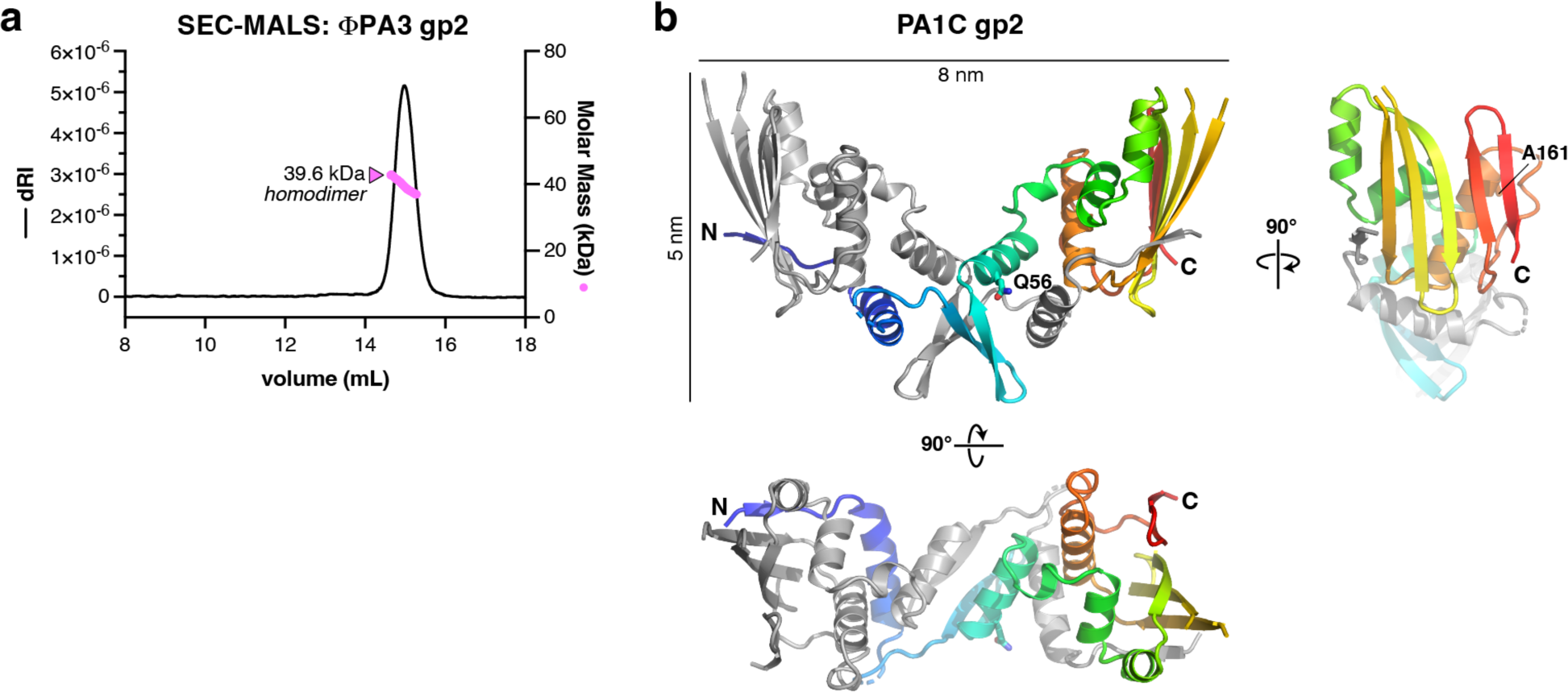
Structure of gp2. (a) Size exclusion chromatography coupled to multi-angle light scattering (SEC-MALS) of ΦPA3 gp2, showing that it is homodimeric in solution (monomer molecular weight = 22.5 kDa). See **Figure S5a-c** for SEC-MALS analysis of other jumbo phage gp2 proteins. (b) Structure of the PA1C gp2 homodimer, with one protomer colored gray and the other colored as a rainbow from N-terminus (blue) to C-terminus (red).

### ChmB point mutants disrupt phage nucleus formation

We next aligned ChmB homologs from jumbo phages that infect *Pseudomonas* species (**Figure S5**) and identified two highly conserved surface residues: Q53 and A159 (ΦPA3 gp2 numbering). To determine the roles of ChmB in phage nucleus formation and function, we expressed GFP-tagged wild-type ΦPA3 ChmB and two point mutants, Q53A and A159D, in ΦPA3-infected *P. aeruginosa* cells (**Figure 4a**). When we overexpressed GFP-tagged wild-type ChmB in infected cells, we observed a striking increase in the average size of the phage nucleus, and a concomitant increase in the nuclear DNA content (as measured by total DAPI signal within the nucleus) (**Figure 4b,c**). Further, a significant fraction of cells (42%) showed ChmB localization suggestive of multiple juxtaposed phage nuclei, or a single phage nucleus with an aberrant shell structure (**Figure 4d**).

**Figure 4.**
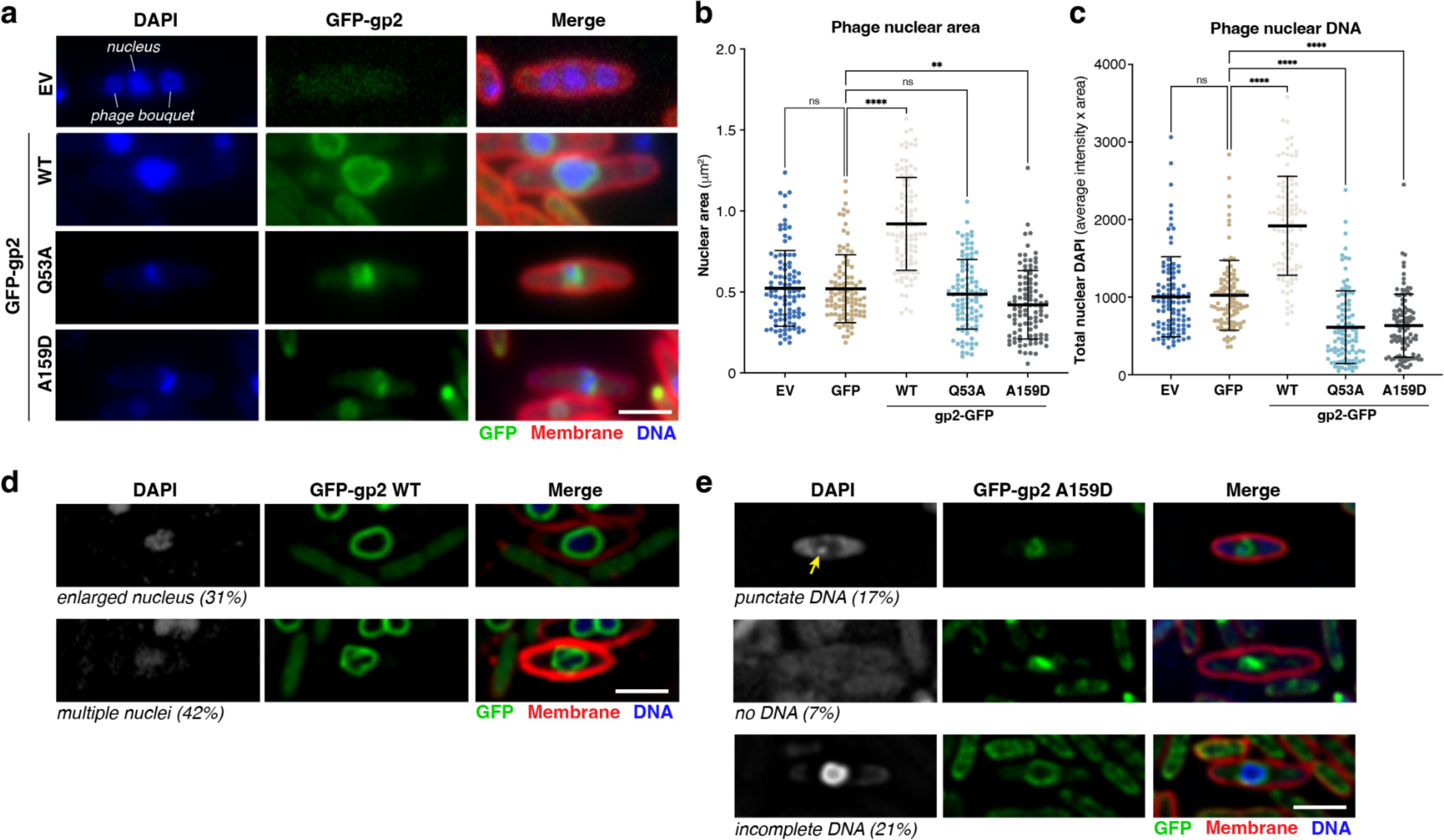
gp2 mutations cause defects in phage nucleus formation and morphology. (a) Fluorescence imaging of ΦPA3-infected *P. aeruginosa* cells expressing no additional proteins (EV: empty vector), GFP-tagged wild-type gp2, or GFP-tagged gp2 point mutants. Un-deconvolved images that were used for DAPI quantitation (panels b and c) are shown. GFP is shown in green, FM4-64 (to visualize membranes) in red, and DAPI (to visualize nucleic acids) in blue. Scale bar = 2 μm. (b) Phage nuclear area of ΦPA3-infected *P. aeruginosa* cells expressing no additional proteins (EV: empty vector), GFP-tagged wild-type gp2, or GFP-tagged gp2 point mutants. n=100 for all samples; error bars represent mean +/- standard deviation. P-values: ns: p>0.05 (not significant); **:p<0.01; ****:p<0.0001. (c) Total nuclear DNA in ΦPA3-infected *P. aeruginosa* cells expressing no additional proteins (EV: empty vector), GFP-tagged wild-type gp2, or GFP-tagged gp2 point mutants, calculated by multiplying each cell’s average DAPI signal within the nucleus by that cell’s nuclear area (panel (b)). (d) Visual phenotypes observed in ΦPA3-infected *P. aeruginosa* cells expressing GFP-tagged wild-type gp2 (n=100 cells). See **Figure S6c** for additional examples. (e) Visual phenotypes observed in ΦPA3-infected *P. aeruginosa* cells expressing GFP-tagged gp2 A159D (n=100 cells). See **Figure S6D** for additional examples, and **Figure S5e** for examples of similar phenotypes from gp2 Q53A.

Expression of either ChmB-Q53A or A159D reversed the large/multiple phage nucleus phenotype observed when overexpressing wild-type ChmB (**Figure 4b,c**) and caused a significant disruption in nucleus formation and/or DNA content in a large fraction of cells. In around 20% of cells (23% for ChmB-Q53A, 17% for ChmB-A159D), the phage nuclear DNA signal in late infections resembled the puncta usually observed in very early infections, prior to significant phage nuclear DNA replication and nucleus growth (**Figure 4e**)^6^. In another population of cells (5% for ChmB-Q53A, 7% for ChmB-A159D), the nuclear DNA appeared to be entirely absent despite these cells sometimes showing ChmB localization reminiscent of a phage nuclear shell (**Figure 4e**). Finally, a third population of cells (14% for ChmB-Q53A, 21% for ChmB-A159D) showed an aberrant nuclear shell structure with nuclear DNA staining that appeared to incompletely fill the nuclear area (**Figure 4e**). Overall, nearly half of infected cells expressing mutant ChmB (42% for ChmB-Q53A, 45% for ChmB-A159D) showed abnormal nuclear shell and/or nuclear DNA morphology. Together, these data show that ChmB is critical for the proper formation and development of the jumbo phage nuclear shell. Further, they imply that disruption of ChmB function affects the replication of genomic DNA, either directly or indirectly by affecting the ability of the phage to properly transport macromolecules into or out of the nucleus.

## Discussion

In contrast to well-studied small-genome phages, nucleus-forming jumbo phages form several characteristic structures in infected cells including the phage nucleus^6, 7, 9, 15^, PhuZ spindle^6, 7, 13, 30^, and phage bouquets^14, 15^. The distinctive subcellular organization imposed by these phages enabled us to use proximity labeling with a phage for the first time, allowing the identification of dozens of phage proteins that localize to particular structures. By combining proximity labeling with subcellular localization analysis by fluorescence microscopy, we established a subcellular protein localization map consisting of 42 phage proteins (**Figure 5a**).

**Figure 5.**
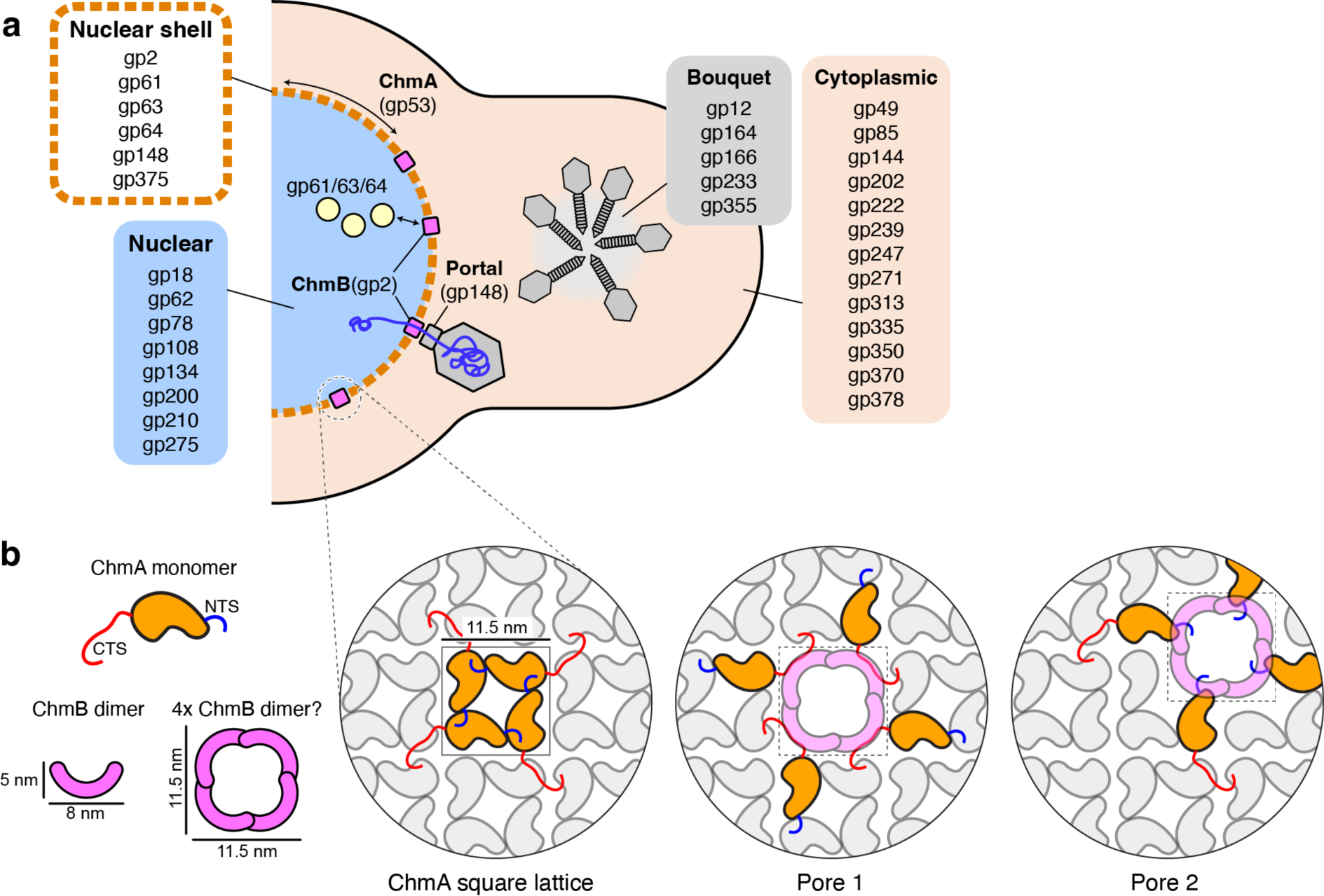
Model for jumbo phage protein localization and nuclear shell architecture and function. (a) Schematic of a ΦPA3-infected *P. aeruginosa* cell with assembled phage nucleus (blue) bounded by a ChmA lattice (orange). Proteins that we find localize to the nucleus, nuclear shell, phage bouquet, or cytoplasm are listed. ChmB (pink) is shown integrated into the ChmA lattice, where it may mediate the docking of phage capsids by binding the portal protein, for genomic packaging. Further interactions with gp61/gp63/gp64 (light yellow) or other shell-associated proteins could accommodate mRNA export or specific protein import. (b) Schematic of the ChmA lattice derived from cryoET analysis of intact 201Φ2-1 and Goslar nuclear shells. ChmA is shown in orange with N-terminal and C-terminal tails shown blue and red, respectively. Removal of four contiguous ChmA protomers from the lattice would leave a cavity ∼11.5 × 11.5 nm (two possibilities shown), which could be filled by an assembly of ChmB to generate a pore.

Here, we focus on the phage nucleus, which separates the replicating phage genome from the host cytoplasm and protects the phage from DNA-targeting immune systems encoded by the host. We previously showed that the jumbo phage nuclear shell is predominantly composed of a single layer of the phage-encoded protein chimallin (ChmA), which assembles into a lattice with pores less than 2 nm in width^10^. Because these pores likely too small for the passage of nucleic acids or proteins, we theorized that the ChmA lattice likely incorporates additional components that mediate mRNA export and selective protein import. Further, based on our prior observation that capsids dock on the phage nuclear shell for genome packaging^5^, the shell must also incorporate components that mediate this docking and enable the passage of genomic DNA through the nuclear shell.

Our proximity labeling and localization mapping approach identified six proteins that associate with the jumbo phage nuclear shell. We find that one of these proteins (gp2) associates directly with self-assembled ChmA *in vitro*, suggesting that it is an integral phage shell protein and prompting us to name this protein ChmB. ChmA self-assembles into a flexible square lattice with the extended NTS and CTS regions binding to neighboring protomers (**Figure 5b**)^10^. NTS-mediated interactions define ChmA homotetramers that measure ∼11.5 × 11.5 nm, while CTS interactions mediate interactions between neighboring tetramers. The shape and dimensions of the ChmB dimer, at ∼5 × 8 nm, suggest a model in which multiple ChmB dimers could line a hole created by the removal of four or more neighboring ChmA protomers from the lattice (two possibilities shown in **Figure 5b**).

In addition to associating directly with ChmA, we find that ChmB is at the center of a large protein-protein interaction network in phage-infected cells. ChmB interacts directly with the phage portal protein, and the portal protein can also localize on its own to the phage nuclear shell in infected cells. Based on these data, we propose that ChmB mediates capsid docking and genome packaging at the nuclear shell. ChmB may also interact with other nuclear shell-associated proteins such as gp61, gp63, and gp64 to mediate specific transport of mRNA and protein through the shell.

Overexpression of wild-type ChmB in infected cells resulted in phage nuclei that were significantly larger than normal, often showing aberrant morphology with multiple (up to 5) distinct phage nuclei in a single cell. Conversely, expression of mutant ChmB proteins (Q53A or A159D) led to the formation of significantly smaller phage nuclei or phage nuclei that appeared to be completely devoid of DNA based on DAPI staining. These data support the idea that ChmB plays fundamental roles in the formation and maturation of the phage nucleus.

ChmB is conserved among all known nucleus-forming jumbo phages that infect *Pseudomonas*, but it is not found in distantly related phage such as *E.coli* phage Goslar^15^ or *Serratia* phage PCH45^9^. Given its apparent role in phage nuclear shell function, why is ChmB not more widely conserved? One possibility is that distant homologs of ChmB might be too divergent to be recognized by sequence-based searches. This possibility is supported by the high sequence divergence of gp2 homologs relative to other proteins like ChmA and PhuZ: across four representative jumbo phages infecting *Pseudomonas* (ΦPA3, PA1C, Phabio, and Psa21), pairwise sequence identities for ChmB homologs average 25%, while ChmA proteins average 51% identity and PhuZ proteins average 46% identity. Alternatively, a different protein might be performing a role equivalent to ChmB in other nucleus-forming jumbo phage families.

In summary, this work has revealed new insights into the organizational principles of nucleus-forming jumbo phages, and into the molecular mechanisms of the phage nucleus in particular. Our results identify a protein interaction network centered around the phage nucleus that will form the basis of future research in this area. We identified a key protein, ChmB, at the center of this network that could play multiple roles as both a pore for macromolecular transport through the nuclear shell, and for capsid docking and genomic DNA packaging. Further work will be required to fully understand the composition of the phage nucleus and the myriad proteins that contribute to its remarkable functions.

## Materials and Methods

### Strains, growth condition and phage preparation

*P. chlororaphis* strains cells (200-B) were cultured on solid Hard Agar (HA). *P. aeruginosa* strains (PA01 and K2733(Pump-knockout)) were cultured on Luria-Bertani (LB) media. Cells were incubated at 30°C overnight. 20 μl of phage high-titer phage lysates of each phage: 201ΦPhi2-1, ΦPhiPA3, and 100uL of OD_600_=0.6 were mixed and incubated for 20 minutes at room temperature. Phage-infected cultures were mixed with 5 mL of HA (for 201ΦPhi2-1) or LB top agar (for phage ΦPhiPA3 and ΦPhiKZ) and poured over a HA or LB plate. The plates were then incubated upside-down at 30°C overnight. The next day, plates were incubated with 5 mL of phage buffer for 5 hours at room temperature. Lysates were collected and centrifuged at 15,000 rpm for 10 minutes. Supernatants were stored at 4°C with 0.01% chloroform.

### Plasmid constructions and bacterial transformation

Genes of interest were PCR-amplified from high-titer phage lysates, then ligated into linearized plasmid backbones using an NEBuilder HiFi DNA Assembly Cloning Kit (New England Biolabs Cat. No. E5520S). Recombinant plasmids were transformed into *E. coli* DH5α and planted on LB containing appropriate antibiotics (25 μg/mL gentamicin sulfate, 100 μg/mL ampicillin, 100 μg/mL spectinomycin). Constructs were confirmed by DNA sequencing and subsequently introduced into indicated organisms of interest and selected on LB supplemented with antibiotics with the same concentrations. Selected overnight cultures were stored in 25% glycerol at -80°C **(Table S6)**.

### Fluorescence microscopy of single cell-infection assay

1.2% agarose pads were prepared on concavity slides. Each pad was supplemented with 0.05%-1% arabinose to induce protein expression. In certain experiments, FM4-64 (1 μg/mL) was used for cell membrane staining and DAPI (1 μg/mL) for nucleoid staining. *P. chlororaphis* and *P. aeruginosa* strains were inoculated on the pads and were grown in a humid chamber at 30°C for 2 hours. For the phage infections, 10 μl of phages (10^8^ pfu/mL) were added on the cells and incubated for an additional 40 minutes in 30°C for infection to proceed. At the desired time points, the pads were sealed with a coverslip and fluorescent microscopy was performed with a DeltaVision Spectris Deconvolution Microscopes (Applied Precision, Issaquah, WA, USA). Cells were imaged for at least 8 stacks in the Z-axis from the middle focal plane with 0.15 μm increments. For time-lapse imaging, 8 points were previously selected and imaged by using the UltimateFocus mode. Images were further processed by the deconvolution algorithm (DeltaVision SoftWoRx Image Analysis Program) and analyzed in Fiji^31^.

### Phage nucleus area and DAPI measurements from fixed cells

1.2% agarose pads were prepared on concavity slides containing 1% arabinose and FM4-64 (1 μg/mL). *P. aeruginosa* strains were inoculated on the pads and grown for 2 hours in humid chamber at 30°C. 75 minutes post infection with 10 μl of phages (10^8^ pfu/mL), 20 μl of fixation mixture (2% of 25% Glutaraldehyde, 82% of 16% PFA and 16% of 1M NaPO_4_ at pH=7.4) was added to each pad and incubated at room temperature for 20 minutes. For high levels of nucleoid staining for quantifications, 20 μl of 20 μg/mL added to the fixed cells and incubated for 20 minutes at room temperature. Pads were sealed with coverslips and fluorescent microscopy was performed as above. Images without deconvolution were analyzed by ImageJ from the DAPI channel. In-built measurement tools were used to calculate the Area and DAPI intensity of the observed nucleoid. Statistical analysis was performed in GraphPad Prism.

### Proximity Labeling with miniTurboID

For proximity labeling, overnight cultures of the strains were grown in LB media with 25 μg/mL gentamicin sulfate. Cultures were then diluted to OD_600_ of 0.1 and supplemented with 500 μM biotin. When cells reached OD_600_=0.5, they were diluted 1:10 into 50 mL total volume in 250 mL flasks and grown in LB supplemented with 0.1% arabinose, 500 μM biotin, 25 μg/mL gentamicin sulfate, and 0.2 mM CaCl_2_. When the cells reached OD_600_=0.3, the cells were infected with phages (MOI 3). At 45 minutes post infection, cultures were collected and centrifuged at 4000 rpm at 4°C. Cell pellets were stored at -80°C for mass spectrometry.

### Mass Spectrometry

To prepare biotinylated samples for IP and mass spectrometry, frozen cell pellets (100 μL) were thawed and resuspended in 100 μL water. Ten μL of resuspended cells were mixed with 200 μL of 6M guanidine-HCl, vortexed, and then subjected to three cycles of incubation at 100°C for 5 minutes, followed by cooling to room temperature. 1.8 mL of pure methanol was added to the boiled cell lysate, vortexed, incubated at -20°C for 20 minutes, then centrifuged at 14K RPM for 10 minutes at 4°C. The tube was inverted and dried to remove any liquid, then the pellet was resuspended in 200 μL of 8 M urea in 0.2 M ammonium bicarbonate. The mixture was incubated for 1 hour at 37°C with constant agitation. Following the incubation, 4 μL of 500 mM TCEP (Tris(2-carboxyethyl) phosphine) and 20 μL 400 mM chloro-acetamide were added. Protein concentration was measured by BCA assay from a 10 μL sample, then 600 μL of 200 mM ammonium bicarbonate was added to bring the urea concentration to 2 M. One μg of sequencing-grade trypsin was added for each 100 μg of protein in the sample, then incubated overnight at 42°C. Following trypsin incubation, 50 μL of 50% formic acid was added (ensuring that the pH drops to 2 using pH test strip), then samples were desalted using C18 solid phase extraction (Waters Sep-Pak C18 12 cc Vac Cartridge, WAT036915) as described by the manufacturer protocol. The peptide concentration of each sample was measured using BCA after resuspension in 1 ml PBS buffer.

For biotin IP, 200 μL of 50% slurry of NeutrAvidin beads (Pierce) was washed three times with PBS, then 1 mg of resuspended peptide solution in PBS was added and incubated 1 hour at room temperature. Beads were washed three times with 2 mL PBS plus 2.5% acetonitrile, washed once in ultrapure water, and excess liquid was carefully removed with a micropipette. Biotinylated peptides were eluted twice with 300 μL of elution buffer (0.2% trifluoroacetic acid, 0.1% formic acid, and 80% acetonitrile in water), with the second elution involving two 5-minute incubations at 100°C. Samples were then dried completely prior to mass spectrometry.

### LC-MS-MS

Trypsin-digested peptides were analyzed by ultra-high pressure liquid chromatography (UPLC) coupled with tandem mass spectroscopy (LC-MS/MS) using nano-spray ionization. The nanospray ionization experiments were performed using a Orbitrap fusion Lumos hybrid mass spectrometer (Thermo) interfaced with nano-scale reversed-phase UPLC (Thermo Dionex UltiMate 3000 RSLC nano System) using a 25 cm, 75-micron ID glass capillary packed with 1.7-µm C18 (130) BEH beads (Waters corporation). Peptides were eluted from the C18 column into the mass spectrometer using a linear gradient (5–80%) of ACN (Acetonitrile) at a flow rate of 375 μl/min for 3 hours. The buffers used to create the ACN gradient were: Buffer A (98% H_2_O, 2% ACN, 0.1% formic acid) and Buffer B (100% ACN, 0.1% formic acid). Mass spectrometer parameters are as follows; an MS1 survey scan using the orbitrap detector (mass range (m/z): 400-1500 (using quadrupole isolation), 120000 resolution setting, spray voltage of 2200 V, Ion transfer tube temperature of 275°C, AGC target of 400000, and maximum injection time of 50 ms) was followed by data dependent scans (top speed for most intense ions, with charge state set to only include +2-5 ions, and 5 second exclusion time, while selecting ions with minimal intensities of 50000 at in which the collision event was carried out in the high energy collision cell (HCD Collision Energy of 30%), and the fragment masses where analyzed in the ion trap mass analyzer (With ion trap scan rate of turbo, first mass m/z was 100, AGC Target 5000 and maximum injection time of 35ms). Protein identification and label free quantification was carried out using Peaks Studio 8.5 (Bioinformatics solutions Inc.) Variable modification at lysine residues of +226.08 amu was used in the peptide sequencing parameters.

### Mass Spectrometry Analysis

For each sample, biotinylated peptides identified by mass spectrometry were divided into host (*P. aeruingosa*) and phage (ΦPA3) peptides. Host peptides that were identified in all samples were used to normalize phage peptide signals across the dataset. Biotinylated phage protein peak areas were calculated by summing the peak areas of each peptide assigned to a given protein. The fold changes of proteins from ChmA (gp53)-miniTurboID and RecA (gp175)-miniTurboID were calculated by comparing to protein peak areas from GFP-miniTurboID samples. The proteins were sorted according to their average normalized fold changes.

The genome sequence of ΦPA3 (NCBI RefSeq NC_028999.1) is misannotated between the coding regions for gp64 (NCBI accession #YP_009217147.1, nucleotides 51942-53267) and gp68 (NCBI accession #YP_009217148.1, nucleotides 58478-60010). We manually annotated this region to identify gp65-66 (which together code for a single protein, separated by an intron spanning nucleotides 55005-55454)^19^ and gp67 for identification by mass spectrometry:

>gp65-66 (NC_028999.1 nucleotides 53811-55004 and 55455-56447)

MYEEHNLRRAVREIHAKLLGHAALDPYYGTTSAARGAMFLSHIGQAPVVEGNEPRRVMTGMEMRYAEYTFDVRLPTDCTILHKVRKYPTGQGYGAIQHNPVTTLIYENYYDEYKTIGVLHVPEYMSFHQDFGYELVKNKEVWESLQPDQMFAKDTVIAQSSTVKSNGLYGMGVNANVAFMSVPGTIEDGFVVSDEFLERMSPRTYTTAVCGAGKKAFFLNMYGDDKIYKPFPDIGEKIREDGVIFAVRDLDDDLAPAEMTPRALRTLDRTFDRAVIGDPGATVKDIKVYWDERQNPSFTPSGMDGQLRKYYDALCTYYREIIKIYRGLLARRKDKLRISEEFNQLLVEAMIYLPQAEGQRKLTRMYRLEQLDEWRVELTYESIKVPGGAYKLTDFHGGKGVVCEVRPKADMPVDEFGNVVDAIIFGGSTMRRSNYGRIYEHGFGAASRDLAQRLRVEAGLPRHGVVPEQDLNRVCSNREWVTYAFAELQEFYYIIAPTMHEILREHPSPAEYVKTVLRDGFSYIYSPVDDPVDLMSSLNCIMNSRFCPNHTRVTYRGQDGKMVTTKDKVLVGPLYMMLLEKIGEDWSAVASVKVQQFGLPSKLNNSDRSSTPGRESAIRSFGESETRSYNCTVGPEATVELLDQTNNPRAHLAVINSILTADKPSNIERAVDRTKVPFGSSRPVDLLEHLLECRGLKFEYATTDGVQPVHTAVPIRAQQKVKSEAIEE*

>gp67 (NC_028999.1 nucleotides 56450-58420)

MNQYNARDLLNMSYDDLFAIPNEWHKIIFDDGEILTKDRATKLSILLWHPLKQFPNATLSVKYHLGDTRVTSKSLVKLLNSVIWGIHAWSNEQVDPEVLARLAIEAKNVLYNEATSRLGAYVATLSMFEIAEVYNHPKVREANQNIEPTTHGIETIAYGKIKEAFNDPTQFRGNSIIEGLRSGTQKMEQLLQAFGPRGFPTDINSDIFAEPCLTGYIDGIWGLYENMIESRSGTKALLYNKELLRVTEYFNRKSQLIAQYVQRLHKGDCGAGYIEFPVIKAYLKSLRGKFYLNEETGKREILQGNETHLIGKKIKMRSVLGCVHPDPQGICATCYGTLADNIPRGTNIGQVSAVSMGDKITSSVLSTKHTDATSAVEQYKITGVEAKYLREGQAPETLYLKKELANKGYRLMIGRNEAQNLADVLMIDNLSAYPPTSASELTRIGLVRTVDGIDEGDVLTVSLYNRKASLSIELLQHVKRVRWELDNRDNIVIDLNGFDFSLPFLTLPYKHVNMYEVMKRIQSFLHSGSDTEGSKLSSDKVGFTSKTYLKNYNDPIDAVAAFASLVNEKIQLPMPHCEVLVYAMMVRSTQQRDYRLPKPGISGQFEKYNKLMQSRSLAGAMAFEKQHEPLNNPGSFLYTLRNDHPYDLAVKGGKLY*

### GFP pulldowns

For GFP pulldowns, GFP-Trap Magnetic Agarose beads (Chromotek gtma-20) were used. Overnight cultures of strains expressing GFP-tagged proteins of interest were grown in LB plus 25 μg/mL gentamicin sulfate. Cells were diluted to an OD_600_ of 0.1, then grown further to an OD_600_ of 0.5. Cultured were diluted 1:10 into 50 mL total volume of culture in 250 mL flasks and grown in LB supplemented with 0.1% arabinose, 25 μg/mL gentamicin sulfate, and 0.2 mM CaCl_2_. When the cells reached an OD_600_ of 0.3, the cells were infected with phages at MOI 3. The cultures were collected at 45 minutes post infection and centrifuged at 4000 rpm at 4°C. Cell pellets were stored at -80°C before the pulldown.

The cell pellets were incubated for 1 hour with 500 μL lysis Buffer (10% glycerol, 25 mM Tris (pH 7.5), 150 mM NaCl, 4 mg/mL lysozyme, 20 μg/mL DNase I, 2x cOmplete Protease Inhibitor, 0.4 mM PMSF). Cell suspensions were sonicated for 10 rounds × 20 pulses/round (Duty Cycle 40, Output 4). Suspension was centrifuged for 30 minutes at 15,000 rpm at 4°C. The 25 μL of bead slurry was used for each sample and equilibrated with Dilution buffer (10 mM Tris-HCl pH 7.5, 150 mM NaCl, 0.5 mM EDTA) and magnetic eppendorf tube rack for 3 times. 500 μL of cell lysate was added to the beads and rotated end-to-end for an hour at 4°C. Cells-bead mixture was washed 5 times with Wash buffer (10 mM Tris/Cl pH 7.5, 150 mM NaCl, 0.05 % Nonidet™ P40 Substitute, 0.5 mM EDTA). At the final wash, samples were transferred to an eppendorf tube. For SDS-Page, cells were resuspended with 2x SDS-buffer (120 mM Tris-HCl pH 6.8, 20% glycerol, 4% SDS, 0.04% bromophenol blue, 10% β-mercaptoethanol) and boiled at 100°C for 5 minutes. 10 μL samples of each elution were run on two separate SDS-PAGE gels and visualized by either silver staining or Coomassie blue staining. For tryptic mass spectrometry of gel bands, bands were cut out Coomassie blue-stained gels. The remaining 80% of each elution was used for mass spectrometry identification of proteins as described above.

### Protein purification

Codon-optimized sequences encoding full-length phage gp2 homologs were synthesized (Invitrogen/GeneArt) and cloned into UC Berkeley Macrolab vector 2BT (Addgene #29666) to generate constructs with N-terminal TEV protease-cleavable His_6_-tags. Proteins were expressed in *E. coli* strain Rosetta 2 (DE3) pLysS (EMD Millipore). Cultures were grown at 37°C to OD_600_=0.7, then induced with 0.25 mM IPTG and shifted to 20°C for 16 hours. Cells were harvested by centrifugation and resuspended in buffer A (25 mM Tris pH 7.5, 10% glycerol, and 1 mM NaN_3_) plus 300 mM NaCl, 5 mM imidazole, 5 mM β-mercaptoethanol. Proteins were purified by Ni^2+^-affinity (Ni-NTA agarose, Qiagen) then passed over an anion-exchange column (Hitrap Q HP, GE Life Sciences) in Buffer A plus 100 mM NaCl and 5 mM β-mercaptoethanol, collecting flow-through fractions. Tags were cleaved with TEV protease^32^, and cleaved protein was passed over another Ni^2+^ column (collecting flow-through fractions) to remove uncleaved protein, cleaved tags, and tagged TEV protease. The protein was passed over a size exclusion column (Superdex 200, GE Life Sciences) in buffer GF (buffer A plus 300 mM NaCl and 1 mM dithiothreitol (DTT)), then concentrated by ultrafiltration (Amicon Ultra, EMD Millipore) to 10 mg/ml and stored at 4°C.

For characterization of oligomeric state by size exclusion chromatography coupled to multi-angle light scattering (SEC-MALS), 100 µL of purified proteins at 5 mg/mL were injected onto a Superdex 200 Increase 10/300 GL column (GE Life Sciences) in buffer GF. Light scattering and refractive index profiles were collected by miniDAWN TREOS and Optilab T-rEX detectors (Wyatt Technology), respectively, and molecular weight was calculated using ASTRA v. 6 software (Wyatt Technology).

### Crystallization and structure determination

Purified PA1C gp2 in a buffer containing 20 mM Tris pH 8.5, 1 mM DTT, and 100 mM NaCl (14 mg/mL) was mixed 1:1 with well solution containing 0.1 M Tris pH 8.5, and 1.5 M Lithium sulfate in hanging drop format. Crystals were cryoprotected by the addition of 24% glycerol and flash-frozen in liquid nitrogen. Diffraction data were collected at the Advanced Photon Source NE-CAT beamline 24ID-C (see support statement below) and processed with the RAPD data-processing pipeline (https://github.com/RAPD/RAPD), which uses XDS^33^ for data indexing and reduction, AIMLESS^34^ for scaling, and TRUNCATE^35^ for conversion to structure factors. We determined the structure by molecular replacement in PHASER^36^, using a predicted structure from AlphaFold2 (as implemented by the ColabFold consortium) as a search model. We manually rebuilt the initial model in COOT^37^, and refined in phenix.refine^38^ using positional and individual B-factor refinement (**Table S5**).

### APS Support Statement

This work is based upon research conducted at the Northeastern Collaborative Access Team beamlines, which are funded by the National Institute of General Medical Sciences from the National Institutes of Health (P30 GM124165). This research used resources of the Advanced Photon Source, a U.S. Department of Energy (DOE) Office of Science User Facility operated for the DOE Office of Science by Argonne National Laboratory under Contract No. DE-AC02-06CH11357.

## Data Availability

Final refined coordinates and reduced diffraction data for the structure of PA1C gp2 is available at the RCSB Protein Data Bank (www.rcsb.org) under accession ID 7UYX. Raw diffraction data for the structure of PA1C gp2 is available at the SBGrid Data Bank (data.sbgrid.org) under accession ID 908.

## Acknowledgements

The authors thank members of the Pogliano and Corbett labs for helpful discussions and for critical reading of the manuscript. The authors acknowledge support from the National Institutes of Health (R01 GM129245 to J.P. and E.V.; R35 GM144121 to K.D.C.; and NIH shared instrumentation grant S10 OD021724), and the Howard Hughes Medical Institute Emerging Pathogens Initiative (to E.V., J.P., and K.D.C.). E.V. is a Howard Hughes Medical Institute Investigator.

## Author Contributions

E.E. conceived of the study and performed all cloning, microscopy, protein purification, and preparation of samples for mass spectrometry, plus data analysis, figure preparation, and manuscript preparation. A.D. performed x-ray crystallography data collection and analysis. Y.G. performed SEC-MALS analysis. K.T.N. and V.C. cloned and characterized GFP fusions of phage proteins. E.A. performed data analysis and provided input on manuscript preparation. M.G. performed mass spectrometry experiments and initial mass spectrometry data analysis. E.V. provided input on experimental design and interpretation. J.P. conceived of the study, guided experimental design and data interpretation, provided funding, prepared figures, and wrote the manuscript. K.D.C. guided experimental design and data interpretation, provided funding, prepared figures, and wrote the manuscript.

## References

1. Salmond, G. P. C. & Fineran, P. C. A century of the phage: past, present and future. Nat. Rev. Microbiol. 13, 777–786 (2015).

2. Serwer, P., Hayes, S. J., Thomas, J. A. & Hardies, S. C. Propagating the missing bacteriophages: a large bacteriophage in a new class. Virol. J. 4, 21 (2007).

3. Al-Shayeb, B., Sachdeva, R., Chen, L.-X., Ward, F., Munk, P., Devoto, A., Castelle, C. J., Olm, M. R., Bouma-Gregson, K., Amano, Y., He, C., Méheust, R., Brooks, B., Thomas, A., Lavy, A., Matheus-Carnevali, P., Sun, C., Goltsman, D. S. A., Borton, M. A., Sharrar, A., Jaffe, A. L., Nelson, T. C., Kantor, R., Keren, R., Lane, K. R., Farag, I. F., Lei, S., Finstad, K., Amundson, R., Anantharaman, K., Zhou, J., Probst, A. J., Power, M. E., Tringe, S. G., Li, W.-J., Wrighton, K., Harrison, S., Morowitz, M., Relman, D. A., Doudna, J. A., Lehours, A.-C., Warren, L., Cate, J. H. D., Santini, J. M. & Banfield, J. F. Clades of huge phages from across Earth’s ecosystems. Nature 578, 425–431 (2020).

4. M Iyer, L., Anantharaman, V., Krishnan, A., Burroughs, A. M. & Aravind, L. Jumbo Phages: A Comparative Genomic Overview of Core Functions and Adaptions for Biological Conflicts. Viruses 13, (2021).

5. Chaikeeratisak, V., Birkholz, E. A. & Pogliano, J. The Phage Nucleus and PhuZ Spindle: Defining Features of the Subcellular Organization and Speciation of Nucleus-Forming Jumbo Phages. Front. Microbiol. 12, 641317 (2021).

6. Chaikeeratisak, V., Nguyen, K., Khanna, K., Brilot, A. F., Erb, M. L., Coker, J. K. C., Vavilina, A., Newton, G. L., Buschauer, R., Pogliano, K., Villa, E., Agard, D. A. & Pogliano, J. Assembly of a nucleus-like structure during viral replication in bacteria. Science 355, 194–197 (2017).

7. Chaikeeratisak, V., Nguyen, K., Egan, M. E., Erb, M. L., Vavilina, A. & Pogliano, J. The Phage Nucleus and Tubulin Spindle Are Conserved among Large Pseudomonas Phages. Cell Rep. 20, 1563–1571 (2017).

8. Mendoza, S. D., Nieweglowska, E. S., Govindarajan, S., Leon, L. M., Berry, J. D., Tiwari, A., Chaikeeratisak, V., Pogliano, J., Agard, D. A. & Bondy-Denomy, J. A bacteriophage nucleus-like compartment shields DNA from CRISPR nucleases. Nature 577, 244–248 (2020).

9. Malone, L. M., Warring, S. L., Jackson, S. A., Warnecke, C., Gardner, P. P., Gumy, L. F. & Fineran, P. C. A jumbo phage that forms a nucleus-like structure evades CRISPR-Cas DNA targeting but is vulnerable to type III RNA-based immunity. Nat Microbiol 5, 48–55 (2020).

10. Laughlin, T. G., Deep, A., Prichard, A. M., Seitz, C., Gu, Y., Enustun, E., Suslov, S., Khanna, K., Birkholz, E. A., Armbruster, E., Andrew McCammon, J., Amaro, R. E., Pogliano, J., Corbett, K. D. & Villa, E. Architecture and self-assembly of the jumbo bacteriophage nuclear shell. bioRxiv 2022.02.14.480162 (2022). doi:10.1101/2022.02.14.480162

11. Nieweglowska, E. S., Brilot, A. F., Méndez-Moran, M., Baek, M., Li, J., Kokontis, C., Cheng, Y., Baker, D., Bondy-Denomy, J. & Agard, D. A. The ϕPA3 Phage Nucleus is enclosed by a self-assembling, 2D crystalline lattice. bioRxiv (2022). doi:10.1101/2022.04.06.487387

12. Nguyen, K. T., Sugie, J., Khanna, K., Egan, M. E., Birkholz, E. A., Lee, J., Beierschmitt, C., Villa, E. & Pogliano, J. Selective transport of fluorescent proteins into the phage nucleus. PLoS One 16, e0251429 (2021).

13. Chaikeeratisak, V., Khanna, K., Nguyen, K. T., Sugie, J., Egan, M. E., Erb, M. L., Vavilina, A., Nonejuie, P., Nieweglowska, E., Pogliano, K., Agard, D. A., Villa, E. & Pogliano, J. Viral Capsid Trafficking along Treadmilling Tubulin Filaments in Bacteria. Cell 177, 1771–1780.e12 (2019).

14. Chaikeeratisak Vorrapon, Khanna Kanika, Nguyen Katrina T., Egan MacKennon E., Enustun Eray, Armbruster Emily, Lee Jina, Pogliano Kit, Villa Elizabeth & Pogliano Joe. Subcellular organization of viral particles during maturation of nucleus-forming jumbo phage. Science Advances 8, eabj9670 (2022).

15. Birkholz, E. A., Laughlin, T. G., Suslov, S., Armbruster, E., Lee, J., Wittmann, J., Corbett, K. D., Villa, E. & Pogliano, J. A Cytoskeletal Vortex Drives Phage Nucleus Rotation During Jumbo Phage Replication in E. coli. bioRxiv 2021.10.25.465362 (2021). doi:10.1101/2021.10.25.465362

16. Branon, T. C., Bosch, J. A., Sanchez, A. D., Udeshi, N. D., Svinkina, T., Carr, S. A., Feldman, J. L., Perrimon, N. & Ting, A. Y. Efficient proximity labeling in living cells and organisms with TurboID. Nat. Biotechnol. 36, 880–887 (2018).

17. Monson, R., Foulds, I., Foweraker, J., Welch, M. & Salmond, G. P. C. The Pseudomonas aeruginosa generalized transducing phage phiPA3 is a new member of the phiKZ-like group of ‘jumbo’ phages, and infects model laboratory strains and clinical isolates from cystic fibrosis patients. Microbiology 157, 859–867 (2011).

18. Thomas, J. A., Weintraub, S. T., Wu, W., Winkler, D. C., Cheng, N., Steven, A. C. & Black, L. W. Extensive proteolysis of head and inner body proteins by a morphogenetic protease in the giant Pseudomonas aeruginosa phage φKZ. Mol. Microbiol. 84, 324–339 (2012).

19. deYMartín Garrido, N., Orekhova, M., Lai Wan Loong, Y. T. E., Litvinova, A., Ramlaul, K., Artamonova, T., Melnikov, A. S., Serdobintsev, P., Aylett, C. H. S. & Yakunina, M. Structure of the bacteriophage PhiKZ non-virion RNA polymerase. Nucleic Acids Res. 49, 7732–7739 (2021).

20. Thomas, J. A., Rolando, M. R., Carroll, C. A., Shen, P. S., Belnap, D. M., Weintraub, S. T., Serwer, P. & Hardies, S. C. Characterization of Pseudomonas chlororaphis myovirus 201φphi2-1 via genomic sequencing, mass spectrometry, and electron microscopy. Virology 376, 330–338 (2008).

21. Thomas, J. A., Weintraub, S. T., Hakala, K., Serwer, P. & Hardies, S. C. Proteome of the large Pseudomonas myovirus 201φ2-1: delineation of proteolytically processed virion proteins. Mol. Cell. Proteomics 9, 940–951 (2010).

22. Reilly, E. R., Abajorga, M. K., Kiser, C., Mohd Redzuan, N. H., Haidar, Z., Adams, L. E., Diaz, R., Pinzon, J. A., Hudson, A. O., Black, L. W., Hsia, R.-C., Weintraub, S. T. & Thomas, J. A. A Cut above the Rest: Characterization of the Assembly of a Large Viral Icosahedral Capsid. Viruses 12, (2020).

23. Sun, L., Zhang, X., Gao, S., Rao, P. A., Padilla-Sanchez, V., Chen, Z., Sun, S., Xiang, Y., Subramaniam, S., Rao, V. B. & Rossmann, M. G. Cryo-EM structure of the bacteriophage T4 portal protein assembly at near-atomic resolution. Nat. Commun. 6, 7548 (2015).

24. Mitchell, M. S. & Rao, V. B. Functional analysis of the bacteriophage T4 DNA-packaging ATPase motor. J. Biol. Chem. 281, 518–527 (2006).

25. Morita, M., Tasaka, M. & Fujisawa, H. Analysis of the fine structure of the prohead binding domain of the packaging protein of bacteriophage T3 using a hexapeptide, an analog of a prohead binding site. Virology 211, 516–524 (1995).

26. Valpuesta, J. M. & Carrascosa, J. L. Structure of viral connectors and their function in bacteriophage assembly and DNA packaging. Q. Rev. Biophys. 27, 107–155 (1994).

27. Yeo, A. & Feiss, M. Specific Interaction of Terminase, the DNA Packaging Enzyme of Bacteriophage γ, with the Portal Protein of theeh Prohead. J. Mol. Biol. 245, 141–150 (1995).

28. Holm, L. Using Dali for Protein Structure Comparison. Methods Mol. Biol. 2112, 29–42 (2020).

29. van Kempen, M., Kim, S. S., Tumescheit, C., Mirdita, M., Söding, J. & Steinegger, M. Foldseek: fast and accurate protein structure search. bioRxiv 2022.02.07.479398 (2022). doi:10.1101/2022.02.07.479398

30. Kraemer, J. A., Erb, M. L., Waddling, C. A., Montabana, E. A., Zehr, E. A., Wang, H., Nguyen, K., Pham, D. S. L., Agard, D. A. & Pogliano, J. A phage tubulin assembles dynamic filaments by an atypical mechanism to center viral DNA within the host cell. Cell 149, 1488–1499 (2012).

31. Schindelin, J., Arganda-Carreras, I., Frise, E., Kaynig, V., Longair, M., Pietzsch, T., Preibisch, S., Rueden, C., Saalfeld, S., Schmid, B., Tinevez, J.-Y., White, D. J., Hartenstein, V., Eliceiri, K., Tomancak, P. & Cardona, A. Fiji: an open-source platform for biological-image analysis. Nat. Methods 9, 676–682 (2012).

32. Tropea, J. E., Cherry, S. & Waugh, D. S. in High Throughput Protein Expression and Purification: Methods and Protocols (ed. Doyle, S. A.) 297–307 (Humana Press, 2009).

33. Kabsch, W. Integration, scaling, space-group assignment and post-refinement. Acta Crystallogr. D Biol. Crystallogr. 66, 133–144 (2010).

34. Evans, P. R. & Murshudov, G. N. How good are my data and what is the resolution? Acta Crystallogr. D Biol. Crystallogr. 69, 1204–1214 (2013).

35. Winn, M. D., Ballard, C. C., Cowtan, K. D., Dodson, E. J., Emsley, P., Evans, P. R., Keegan, R. M., Krissinel, E. B., Leslie, A. G. W., McCoy, A. & Others. Overview of the CCP4 suite and current developments. Acta Crystallogr. D Biol. Crystallogr. 67, 235–242 (2011).

36. McCoy, A. J., Grosse-Kunstleve, R. W., Adams, P. D., Winn, M. D., Storoni, L. C. & Read, R. J. Phaser crystallographic software. J. Appl. Crystallogr. 40, 658–674 (2007).

37. Emsley, P., Lohkamp, B., Scott, W. G. & Cowtan, K. Features and development of Coot. Acta Crystallogr. D Biol. Crystallog*r.* 66, 486–501 (2010).

38. Afonine, P. V., Grosse-Kunstleve, R. W., Echols, N., Headd, J. J., Moriarty, N. W., Mustyakimov, M., Terwilliger, T. C., Urzhumtsev, A., Zwart, P. H. & Adams, P. D. Towards automated crystallographic structure refinement with phenix.refine. Acta Crystallogr. D Biol. Crystallogr. 68, 352–367 (2012).

