## Supplemental Figures for "Identification of the bacteriophage nucleus protein interaction network"

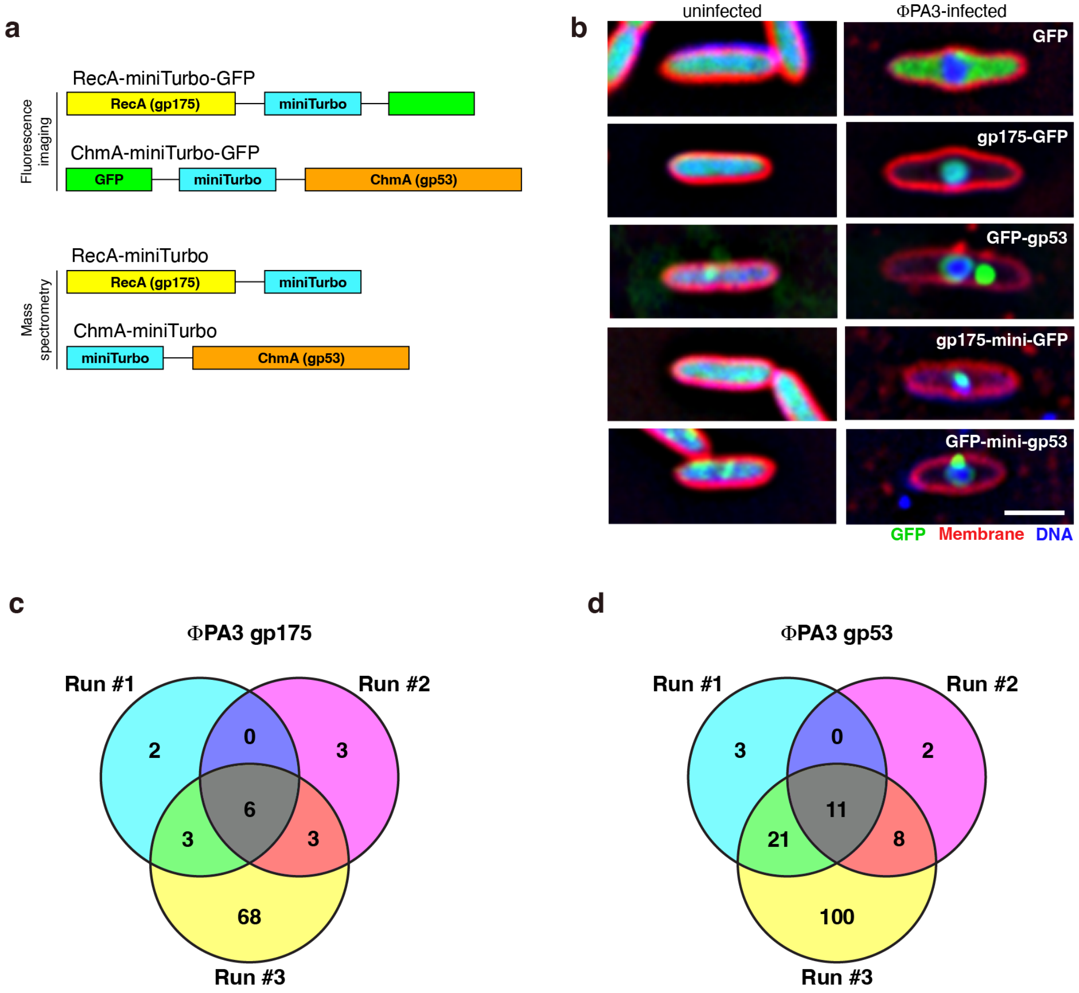


**Figure S1. miniTurboID proximity labeling of phage nucleus-associated proteins**

(a) Construct design for localization and proximity labeling of ΦPA3 RecA (gp175) and ChmA (gp53) associated proteins. (b) Localization of GFP control and GFP- and GFP-miniTurboID tagged RecA (gp175) and ChmA (gp53) in ΦPA3-infected *P. aeruginosa* cells. GFP is shown in green, FM4-64 (to visualize membranes) in red, and DAPI (to visualize nucleic acids) in blue. Scale bar = 2 µm. (c-d) Venn diagrams showing RecA (panel c) and ChmA (panel d) interacting proteins identified by miniTurboID labeling in three independent runs. See **Tables 1-2** and **Tables S1-S2**.

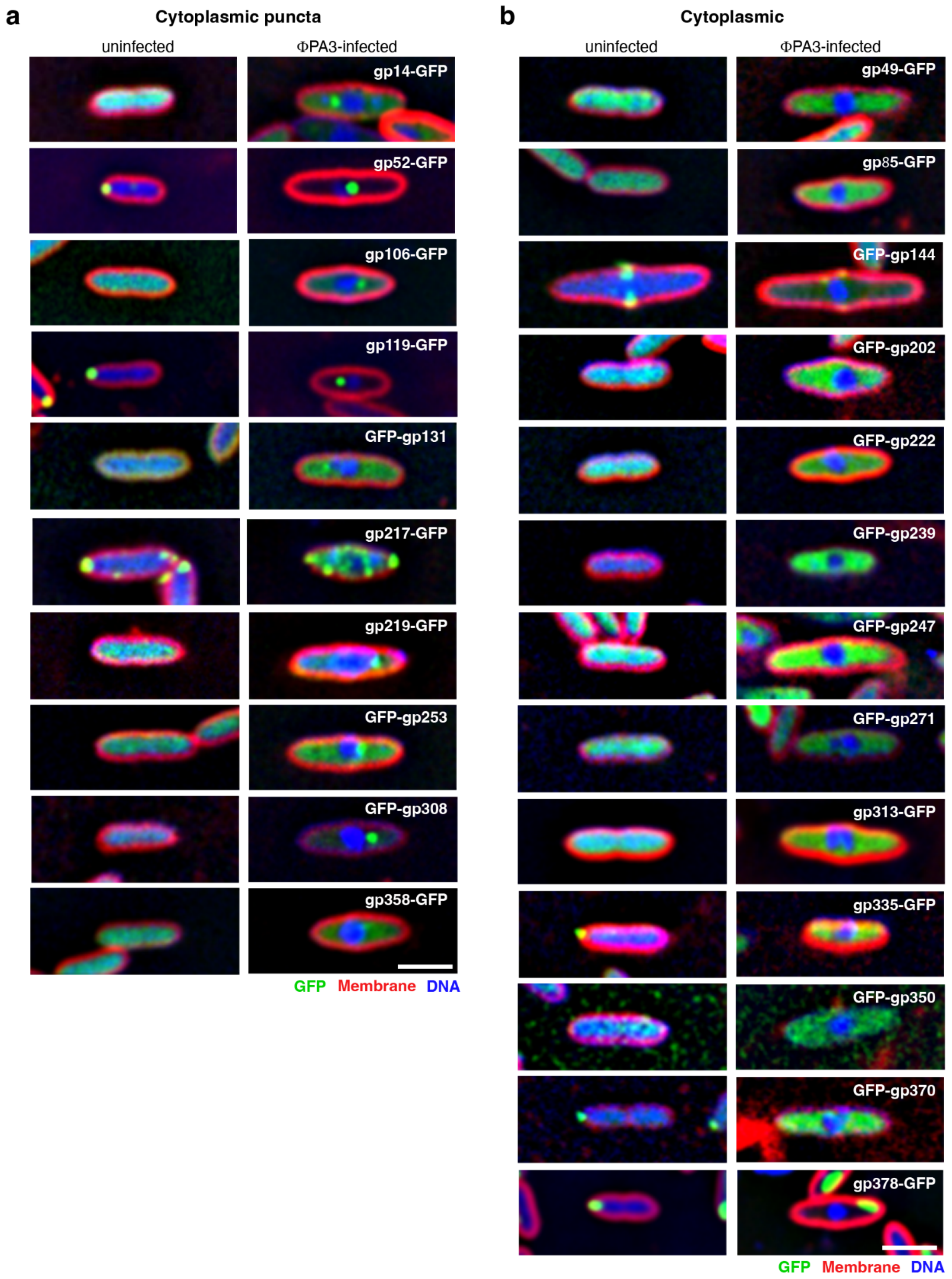


**Figure S2. Localization analysis of RecA- and ChmA-interacting ΦPA3 proteins**

Subcellular localization of selected proteins identified by proximity labeling, with panel (a) showing proteins that localize as cytoplasmic puncta, and (b) showing proteins with diffuse cytoplasmic localization. See **Table 3** for a collated list of localizations. GFP is shown in green, FM4-64 (to visualize membranes) in red, and DAPI (to visualize nucleic acids) in blue. Scale bar = 2 µm.

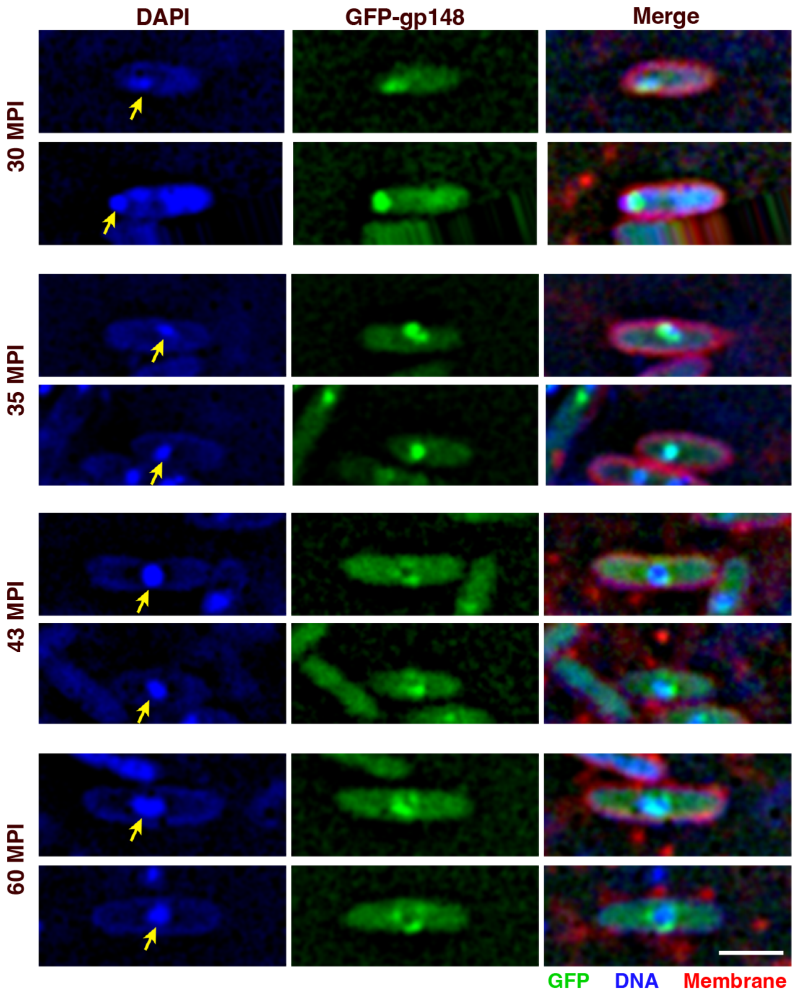


**Figure S3. Early localization of gp148 to the phage nuclear shell**

Subcellular localization of GFP-gp148 in *P. aeruginosa* cells infected with phage ΦPA3, at the indicated times post infection (MPI: minutes post infection). Yellow arrows indicate the position of the phage nucleus. GFP is shown in green, FM4-64 (to visualize membranes) in red, and DAPI (to visualize nucleic acids) in blue. Scale bar = 2 μm.

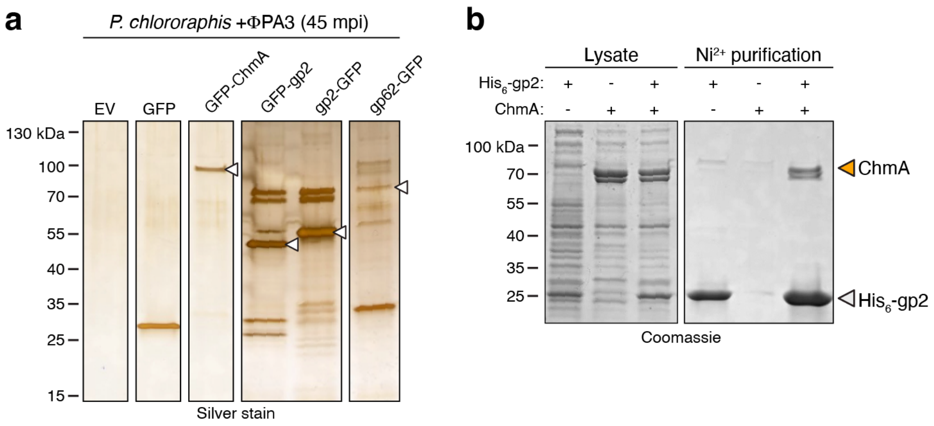


**Figure S4. Protein-protein interaction analysis**

(a) Silver stained SDS-PAGE gels showing GFP pulldown results from GFP-tagged proteins expressed in ΦPA3-infected *P. aeruginosa* (45 minutes post infection). White arrowheads indicate the bait protein for each sample. (b) Coomassie blue-stained SDS-PAGE gel showing $Ni^{2+}$ pulldown results from coexpression of 201Φ2-1 gp2 ($His_6$-tagged) and ChmA (gp105; untagged). ChmA appears as a doublet because of a second start codon at codon 33 of the gp105 gene.

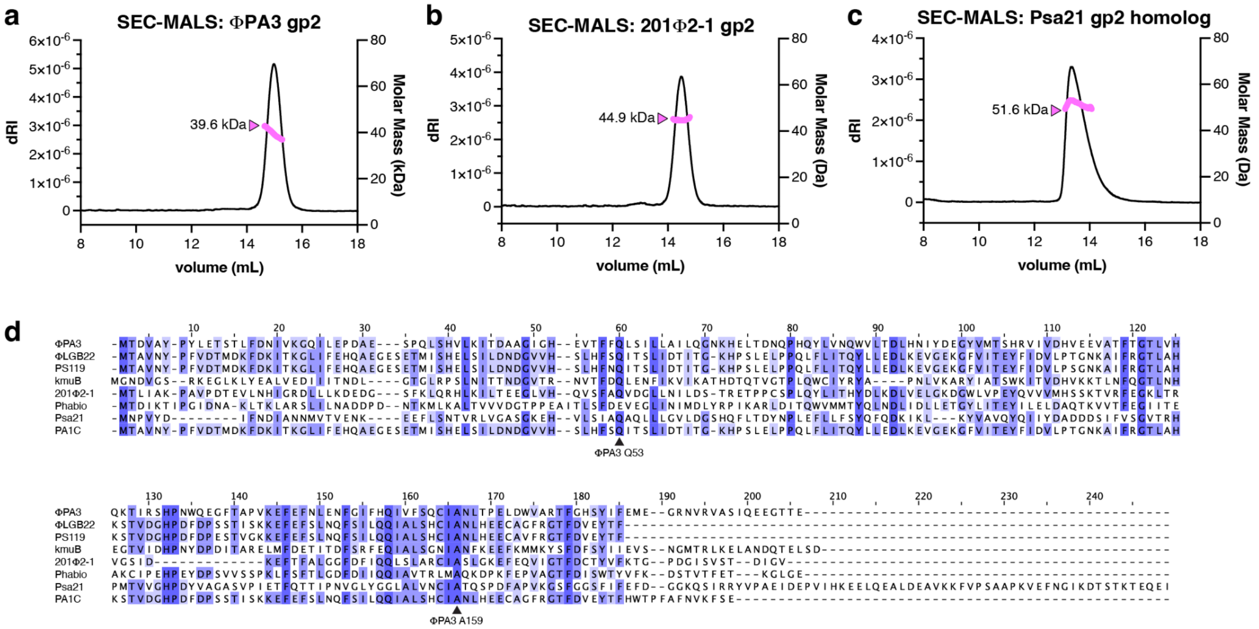

**Figure S5. Biochemical and sequence analysis of jumbo phage gp2 proteins**

(a) Size exclusion chromatography coupled to multi-angle light scattering (SEC-MALS) analysis of ΦPA3 gp2. Measured molecular weight = 39.6 kDa; dimer molecular weight = 45 kDa. (b) SEC-MALS analysis of 201Φ201 gp2. Measured molecular weight = 44.9 kDa; dimer molecular weight = 39.8 kDa. (c) SEC-MALS analysis of the gp2 homolog in phage Psa21 (gp2). Measured molecular weight = 51.6 kDa; dimer molecular weight = 50.6 kDa. (d) Sequence alignment of gp2 homologs in jumbo phage infecting Pseudomonas, showing the position of the highly-conserved Q53 and A159 (ΦPA3 numbering) residues.

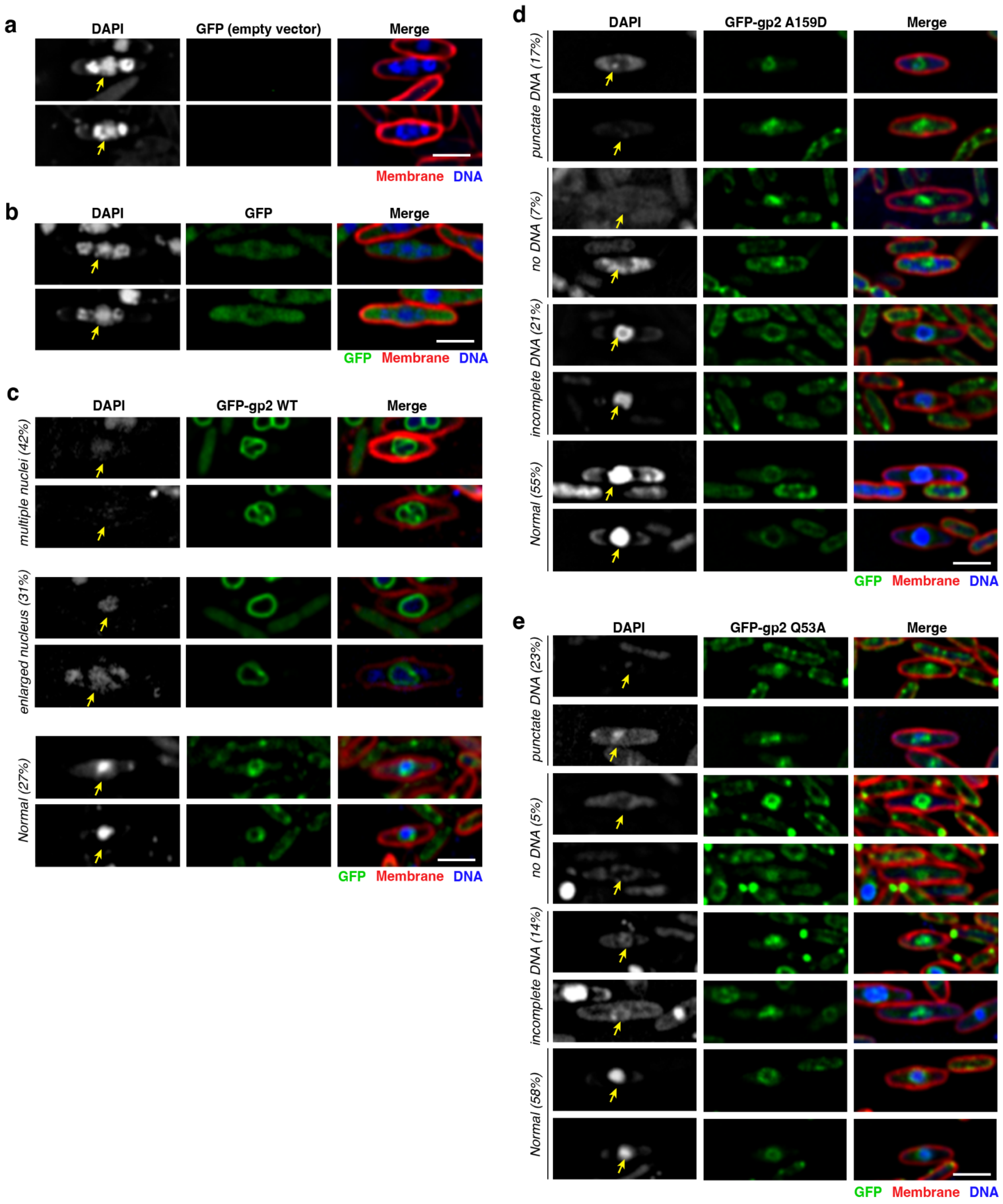


**Figure S6. Overexpression of mutant gp2 proteins affect nuclear shell formation and morphology**

(a) Microscopy of ΦPA3-infected *P. aeruginosa* cells transformed with empty vector. Yellow arrow indicates the position of the phage nucleus. The two DAPI staining bodies bracketing the phage nucleus are the phage bouquets. GFP is shown in green, FM4-64 (to visualize membranes) in red, and DAPI (to visualize nucleic acids) in blue. Scale bar = 2 μm. (b) Microscopy of ΦPA3-infected *P. aeruginosa* cells expressing GFP. (c) Microscopy of ΦPA3-infected *P. aeruginosa* cells expressing GFP-tagged wild-type gp2. Common phenotypes and their percentages in the analyzed sample (n=100 cells) are shown. (d) Microscopy of ΦPA3-infected *P. aeruginosa* cells expressing GFP-tagged gp2 A159D. Common phenotypes and their percentages in the analyzed sample (n=100 cells) are shown. (e) Microscopy of ΦPA3-infected *P. aeruginosa* cells expressing GFP-tagged gp2 Q53A. Common phenotypes and their percentages in the analyzed sample (n=100 cells) are shown.
